# Phylogenetic effective sample size

**DOI:** 10.1101/023242

**Authors:** Krzysztof Bartoszek

## Abstract

In this paper I address the question — *how large is a phylogenetic sample?* I propose a definition of a phylogenetic effective sample size for Brownian motion and Ornstein–Uhlenbeck processes — the *regression effective sample size*. I discuss how mutual information can be used to define an effective sample size in the non-normal process case and compare these two definitions to an already present concept of effective sample size (the mean effective sample size). Through a simulation study I find that the AIC_*c*_ is robust if one corrects for the number of species or effective number of species. Lastly I discuss how the concept of the phylogenetic effective sample size can be useful for biodiversity quantification, identification of interesting clades and deciding on the importance of phylogenetic correlations.

## 1 Introduction

One of the reasons to introduce phylogenetic comparative methods (PCMs) in the words of Martins and Hansen [1996], was to address the problem of statistical dependence. They called the issue the “degrees of freedom” or “effective sample size” problem. If we have *n* species related by a phylogenetic tree, unless it is a star phylogeny, then our effective sample size is less than *n* (in extreme cases even one). Taking into consideration the number of independent observations is important in evaluating the accuracy of parameter estimation or hypothesis tests. The performance of such statistical procedures depends on the number of independent data points and not on the observed number of data points [Martins and Hansen, 1996]. Ignoring the correlations (and hence inflating the sample size) results in too narrow confidence intervals, inflated p–values and power. All of this leads to type I and II errors of which the user may be oblivious of.

In a phylogenetic context the calculation of the effective number of observations has not been often addressed directly. In statistical literature effective sample size (ESS) is usually parameter specific, it can be understood as “the number of independent measurements one would need to reach the same amount of information about a parameter as in the original data” [Faes et al., 2009] — in other words how many independent points do we have for estimating a particular parameter. Nunn [p. 145 2011] points out that often phylogenetic comparative methods have been viewed in a restricted manner as a “degrees of freedom” correction procedure that “reduce the number of data points”, due to the nonindependence. Most phylogenetic comparative methods work in the following way — one assumes a model and maximizes the likelihood under that model. Hence, the issue of ESS, as mentioned above, has been taken care of but only for the estimation problem. In other situations, as Nunn [2011] following Pagel [1993] reminds, the “degrees of freedom analogy can be misleading”. It is more important how the variance is partitioned among species. In fact in the case of model selection, or when one wants to know how many “independent” taxa one has e.g. for conservation purposes the situation becomes much more complex. As we will see it is more important how the covariance is structured.

Smith [1994] directly approached the problem of effective sample size. He studied interspecies phenotypic data by a nested ANOVA and *“Determination of the taxonomic levels that account for most of the variation can be used to select a single level at which it is most reasonable to consider the data points as independent*”. From the perspective of modern phylogenetic comparative methods this is a “hack”, as Smith [1994] himself wrote “the method improves the nonindependence problem but does not eliminate it”. From our perspective his work is important, as from the nested ANOVA setup, he partitioned the variance into components from different levels of the phylogeny and then defined the effective sample size as

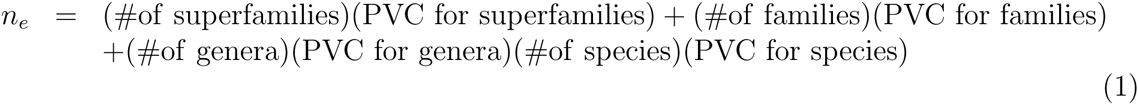

where PVC is percentage of variance component. Smith [1994] importantly notices, that in principle “ *The method does not require that levels of the nested hierarchy are defined by taxonomic categories*.” In this work I develop the idea described in Smith [1994]’s own words: to *“consider each species as some fraction of a free observation varying between* 0 *and* 1.0, *a value could be computed … that would reflect the balance between constraint and independent evolution. This value is defined as the effective sample size (effective N) for the data set and trait, as opposed to the traditionally used observed sample size (observed N).”* Building up on the modern development of stochastic models for phylogenetic comparative methods, I do not have to restrict myself to partitioning the data into hierarchical levels containing different fractions of the variance, but rather look holistically at the dependence pattern induced by the tree and model of evolution.This might make it impossible (but maybe not always) to assign to each species (or taxonomic level) its fraction of free observations but as we shall see it will allow me to calculate the sum of fractions of free observations.

An analysis of phylogenetically structured phenotypic data often has as its goal to identify the mode of evolution, i.e. is the trait(s) adapting (and if so to what trait/phenotype) or rather exhibiting neutral evolution. Information criteria like the Akaike Information Criterion [AIC Akaike, 1974], Akaike Information Criterion corrected for small sample size [AIC_*c*_ Hurvich and Tsai, 1989] or Bayesian Information Criterion [BIC Schwarz, 1978] are commonly used to identify the model better supported by the data. However, if one goes back to the derivation of the AIC_*c*_ [Hurvich and Tsai, 1989] and BIC [Schwarz, 1978] one can see that the *n* observations are assumed independent. Therefore a phylogenetic comparative model seems to violate this assumption, in the best case by inflating the sample size. In a way such an inflation corresponds to not penalizing enough for additional parameters. However in their original paper Hurvich and Tsai [1989] derive the same AIC_*c*_ formula for autoregressive models so this warrants further study in the phylogenetic setting where the covariance structure is hierarchical.

Therefore, using the number of species (unless the phylogeny is a star) results in a risk of overfitting for small phylogenies or those with most speciation events near the tips. In this work I propose a way of taking into account the *effective number of species* during the model selection procedure. The newest version of mvSLOUCH (available fromhttp://cran.r-project.org/web/packages/mvSLOUCH/index.html) allows for automatic model selection if one treats *n* as the true sample size and also if one corrects for the dependencies using an effective sample size. Importantly mvSLOUCH allows for an arbitrary pattern of missing data — no observation is removed and the likelihood is based on all provided information. Using this new version of mvSLOUCH, I include in this work a simulation study and analyze a number of data sets to see how much a difference does it make whether, one uses the observed or effective number of species for model selection. In most cases, the two ways of counting species lead to the same conclusion. However, for small samples (see Tab. 3) using the effective number of species can result in a different outcome. In fact we should expect this to be so, a good correction method should be robust — with enough observations the data (or rather likelihood) should decide no matter how one corrects. It is only with few observations (and hence little power) that correction methods should play a role by pointing to different possibilities of interpreting the observed data.

## 2 Effective sample size

Effective sample size is intuitively meant to represent the number of independent particles of data in the sample. If the sample is correlated, then each observation will only have a certain fraction of the information it carries particular to itself. The rest of the information will be shared with one/some/all other points in the sample. We would like to quantify what proportion of the whole sample is made up of these independent bits of information. If this proportion is p, then our phylogenetic effective sample size (pESS) will be *n*_*e*_ = *pn*. However our situation is a bit different. It is reasonable to assume that we have a least one observation — at least one species described by at least a single trait. One way is to define *p* to be between 1 and 1/*n*. Alternatively we can define as

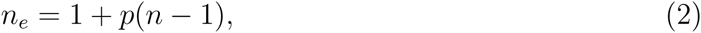

where *p* ∈ [0,1]. I will call this *p* of Eq. (2) the phylogenetic ESS factor. The value *n*_*e*_/*n* is useful in practice to compare between different sized phylogenies and I will call it the relative phylogenetic ESS.

Martins and Hansen [1996] point out, that in the discrete trait case, the ESS cannot be greater than the number of independent evolutionary changes regardless of the number of observed species. Maddison and FitzJohn [2015] very recently remind us of this again. Phylogenetic comparative methods are there to take care of “pseudoreplicates” due to the tree induced correlations. However, especially in the discrete case, tests of significance might have inflated power as one uses the number of species instead of the (unknown) number of independent evolutionary changes. Unfortunately, at the moment, there does not seem to be any solution for this problem [Maddison and FitzJohn, 2015]. Hopefully the phylogenetic effective sample size concept presented here could indicate a direction for finding one. An alternative potential approach in the discrete case, is phylogenetic informativeness based on the number of mutations (i.e. changes) shared by tip taxa under the Poisson process [Mulder and Crawford, 2015, Townsend, 2007]. It however, remains to study the probabilistic properties of phylogenetic informativeness in order to understand whether and how it may be applied in the pESS context.

Statistical definitions of effective sample size are commonly introduced in the context of parameter estimations — what is the ESS for a given parameter/set of parameters. I am in a different situation — I want to quantify how many independent particles do I observe. In this situation one has to propose one’s own definition of effective sample size that will be useful from a practical point of view. This is not an obvious task in the situation of *n* dependent observations. The case of multivariate observations, where individual components are dependent between each other and correlations between traits can be negative, will be even more complicated. Below I will discuss a couple of possible approaches for defining an effective sample size and in the next section discuss how they can be applied in the phylogenetic comparative methods field.

Ané [2008] defined an effective sample size for estimating the root state under a Brownian motion (BM) model of evolution. She noticed that it can be very small — 6 for a phylogeny of 49 species [mammal phylogeny of Garland, T., Jr. et al., 1993]. In fact my simulations and reanalysis of this data (Tab. 3) give very similar numbers. She defined the effective sample size as

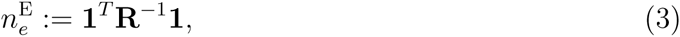

where **R** is the between species correlation matrix. I call 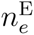 the *mean effective sample size* (mESS), as 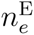 is the number of independent random variables that result in the same precision for estimating the mean value (intercept) of a linear with *n* correlated, by **R**, observations [Ané, 2008]. It is important for the reader to notice that 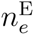 is not connected to any average of sample sizes. The word “mean” in the name refers to the fact that 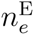 quantifies the information available on the mean value in a linear model.

For our purpose the mean effective sample size is not completely satisfactory. The 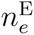 value does not say how much independent signal there is in the sample, but only how much information we have about the expected value. In the scope of this work we are more interested in the former and not the latter. In fact we can observe (Fig. 2 and Tab. 3), that in a phylogenetic sample 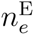 is usually rather low. Such small numbers are due to the high variance of the sample average [an estimator of the mean value Ané, 2008, Bartoszek and Sagitov, 2015b, Sagitov and Bartoszek, 2012], resulting in low precision for the mean value. I therefore consider alternative approaches to define a phylogenetic effective sample size.

**Figure 1:**
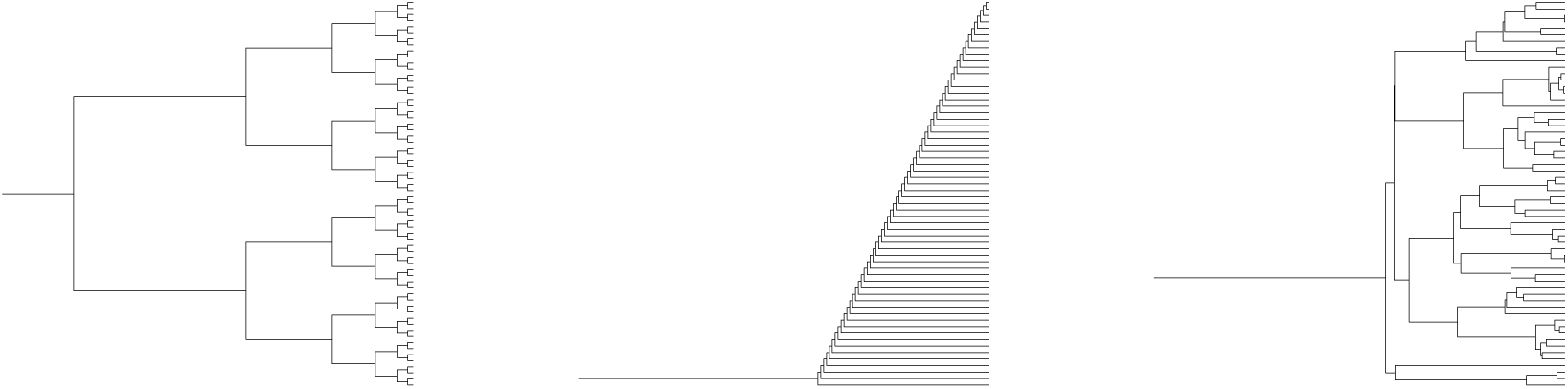
Different binary phylogenetic tree setups used in the simulation studies. Left: fully balanced tree, centre fully unbalanced tree, right: single realization of a pure birth tree. The balanced tree has 64 tips, the other two 60.

**Figure 2:**
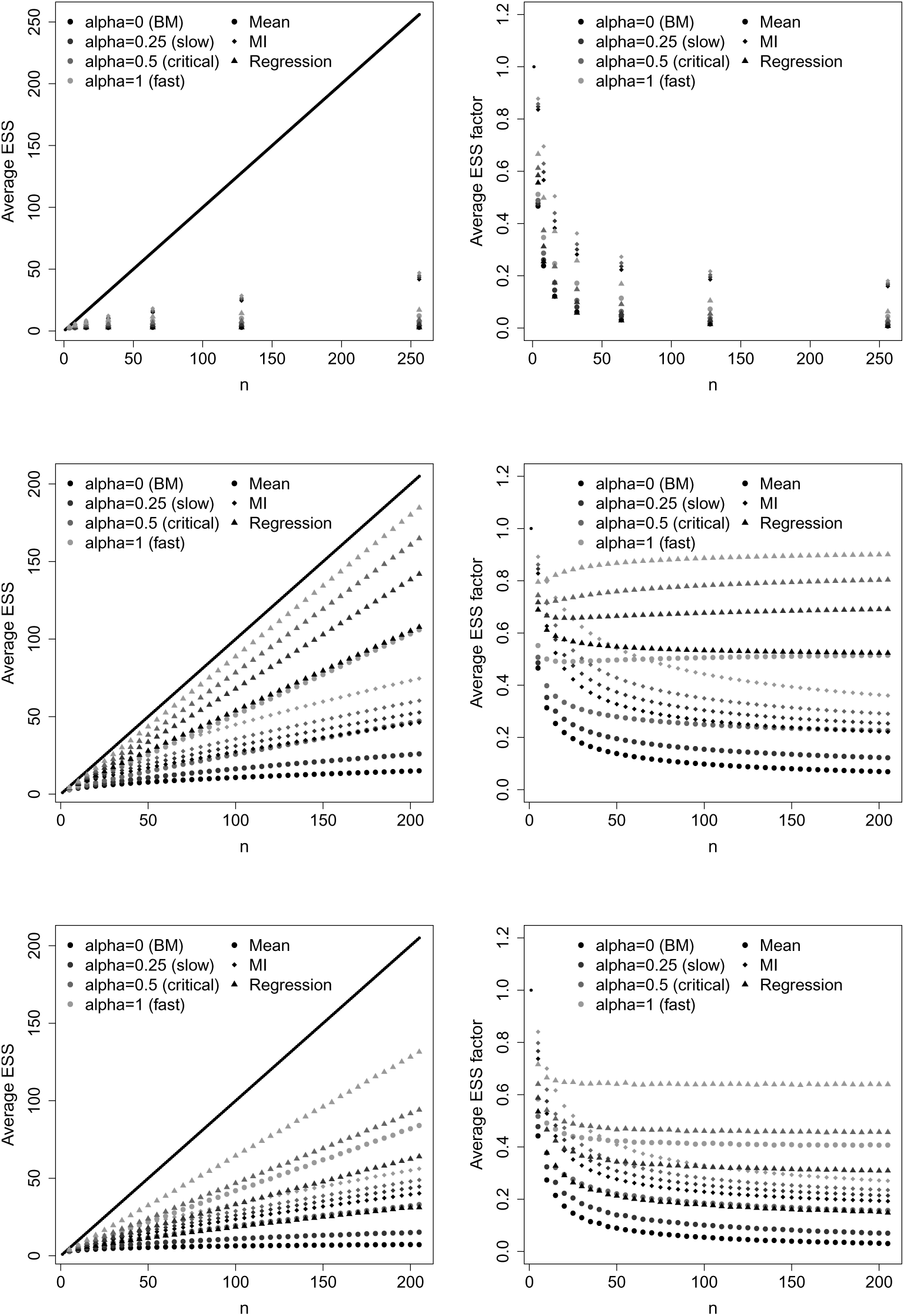
Phylogenetic effective sample sizes for different types of trees and evolutionary processes. First row: balanced tree, second row: left unbalanced tree, third row: average of 1000 pure–birth Yule trees (*λ* = 1). The balanced trees and unbalanced trees were generated using the function stree() of the R [R Core Team, 2013] ape package [Paradis, 2012], the Yule trees by the TreeSim R package [Stadler, 2014, 2011]. First column: phylogenetic effective sample sizes, *n*_*e*_ second column: phylogenetic effective sample size factors, *p*. The parameters of the processes are Brownian motion (*X*_0_ = 0, *σ*^2^ = 1), second row: Ornstein–Uhlenbeck process (*α* = 0.25, *σ*^2^ = 1, *X*_0_ = 0, *θ* = 0), third row: Ornstein–Uhlenbeck process (*σ* = 0.5, *σ*^2^ = 1, *X*_0_ = 0, *θ* = 0), fourth row: Ornstein–Uhlenbeck process (*α* = 1, *σ*^2^ = 1, *X*_0_ = 0, *θ* = 0). The straight black line is the observed number of taxa *n*.

**Table 1:** Comparison of phylogenetic diversity with the proposed pESS definitions for different evolutionary models and topologies. The values are means from a 1000 simulations. The value of 1 for the regression ESS indicates that the calculated value was below 1 and hence the rounding up.

| Model | $n^1$ | E [PD] <sup>2</sup> | E [$n_e^{\text{MI}}$] <sup>3</sup> | E [$n_e^{\text{E}}$] <sup>4</sup> | E [$n_e^{\text{R}}$] <sup>5</sup> |
| --- | --- | --- | --- | --- | --- |
| Yule |  |  |  |  |  |
| BM<sup>6</sup> | 15 | 15.102 | 7.119 | 3.938 | 5.505 |
| slow OU<sup>7</sup> | 15 | 15.102 | 7.744 | 4.661 | 6.881 |
| medium OU<sup>8</sup> | 15 | 15.102 | 8.404 | 5.562 | 8.179 |
| fast OU<sup>9</sup> | 15 | 15.102 | 9.611 | 7.507 | 10.126 |
| unbalanced |  |  |  |  |  |
| BM | 15 | 15.461 | 8.335 | 4.55 | 9.228 |
| slow OU | 15 | 15.461 | 9.045 | 5.208 | 10.217 |
| medium OU | 15 | 15.461 | 9.808 | 6.006 | 11.117 |
| fast OU | 15 | 15.461 | 11.292 | 7.885 | 12.506 |
| balanced |  |  |  |  |  |
| BM | 16 | 8.584 | 6.735 | 2.824 | 2.793 |
| slow OU | 16 | 8.584 | 7.146 | 3.172 | 3.6 |
| medium OU | 16 | 8.584 | 7.598 | 3.608 | 4.532 |

<sup>1</sup> number of tips <sup>2</sup> expected phylogenetic diversity <sup>3</sup> expected mutual information effective sample size <sup>4</sup> expected mean effective sample size <sup>5</sup> expected regression effective sample size <sup>6</sup> Brownian motion $\alpha = 0$ <sup>7</sup> Ornstein–Uhlenbeck $\alpha = 0.25$ <sup>8</sup> Ornstein–Uhlenbeck $\alpha = 0.5$ <sup>9</sup> Ornstein–Uhlenbeck $\alpha = 1$

|  |  |  |  |  |  |
| --- | --- | --- | --- | --- | --- |
| fast OU | 16 | 8.584 | 8.568 | 4.698 | 6.544 |
| balanced short terminal |  |  |  |  |  |
| BM | 16 | 10.829 | 5.892 | 2.689 | 1.294 |
| slow OU | 16 | 10.829 | 6.2 | 3.159 | 1.539 |
| medium OU | 16 | 10.829 | 6.497 | 3.762 | 1.853 |
| fast OU | 16 | 10.829 | 7.003 | 5.078 | 2.57 |
| balanced long terminal |  |  |  |  |  |
| BM | 16 | 44.023 | 15.991 | 15.238 | 15.994 |
| slow OU | 16 | 44.023 | 15.998 | 15.637 | 15.999 |
| medium OU | 16 | 44.023 | 16 | 15.852 | 16 |
| fast OU | 16 | 44.023 | 16 | 15.982 | 16 |
| balanced harmonic decrease |  |  |  |  |  |
| BM | 16 | 19.908 | 7.292 | 3.053 | 4.094 |
| slow OU | 16 | 19.908 | 8.578 | 4.13 | 6.839 |
| medium OU | 16 | 19.908 | 10.107 | 5.694 | 9.634 |
| fast OU | 16 | 19.908 | 12.896 | 9.158 | 13.333 |
| balanced harmonic increase |  |  |  |  |  |
| BM | 16 | 39.702 | 11.242 | 6.25 | 11.196 |
| slow OU | 16 | 39.702 | 13.921 | 9.187 | 14.418 |
| medium OU | 16 | 39.702 | 15.454 | 12.325 | 15.622 |
| fast OU | 16 | 39.702 | 15.986 | 15.379 | 15.987 |
| balanced geometric decrease |  |  |  |  |  |
| BM | 16 | 15.171 | 6.735 | 2.824 | 2.794 |
| slow OU | 16 | 15.171 | 7.54 | 3.549 | 4.41 |
| medium OU | 16 | 15.171 | 8.445 | 4.55 | 6.294 |
| fast OU | 16 | 15.171 | 10.247 | 6.827 | 9.644 |
| balanced geometric increase |  |  |  |  |  |
| BM | 16 | 38.661 | 12.072 | 7.5 | 12.463 |
| slow OU | 16 | 38.661 | 14.335 | 10.198 | 14.75 |
| medium OU | 16 | 38.661 | 15.544 | 12.836 | 15.684 |
| fast OU | 16 | 38.661 | 15.983 | 15.418 | 15.988 |
| balanced long root branch |  |  |  |  |  |
| BM | 16 | 3.598 | 7.045 | 2.824 | 1.914 |
| slow OU | 16 | 3.598 | 7.116 | 2.868 | 2.025 |
| medium OU | 16 | 3.598 | 7.189 | 2.914 | 2.141 |
| fast OU | 16 | 3.598 | 7.34 | 3.011 | 2.382 |
| Yule |  |  |  |  |  |
| BM | 125 | 124.839 | 27.745 | 6.636 | 21.864 |
| slow OU | 125 | 124.839 | 30.748 | 12.229 | 40.549 |
| medium OU | 125 | 124.839 | 33.829 | 22.896 | 58.002 |
| fast OU | 125 | 124.839 | 39.212 | 51.921 | 80.243 |
| unbalanced |  |  |  |  |  |
| BM | 125 | 244.124 | 32 | 11.905 | 66.687 |
| slow OU | 125 | 244.124 | 36.518 | 18.825 | 85.375 |
| medium OU | 125 | 244.124 | 41.85 | 30.96 | 98.777 |
| fast OU | 125 | 244.124 | 52.44 | 64.035 | 111.456 |
| balanced |  |  |  |  |  |
| BM | 128 | 28.19 | 24.599 | 2.977 | 2.796 |
| slow OU | 128 | 28.19 | 25.746 | 4.073 | 5.008 |
| medium OU | 128 | 28.19 | 26.798 | 5.765 | 7.797 |
| fast OU | 128 | 28.19 | 28.526 | 10.189 | 14.288 |
| balanced short terminal |  |  |  |  |  |
| BM | 128 | 36.485 | 24.776 | 2.983 | 3.232 |
| slow OU | 128 | 36.485 | 26.3 | 4.478 | 6.846 |
| medium OU | 128 | 36.485 | 27.662 | 6.886 | 11.571 |
| fast OU | 128 | 36.485 | 29.87 | 12.816 | 21.3 |
| balanced long terminal |  |  |  |  |  |
| BM | 128 | 615.155 | 124.514 | 89.930 | 127.722 |
| slow OU | 128 | 615.155 | 127.759 | 116.316 | 127.980 |
| medium OU | 128 | 615.155 | 127.993 | 125.927 | 127.999 |
| fast OU | 128 | 615.155 | 128 | 127.967 | 128 |
| balanced harmonic decrease |  |  |  |  |  |
| BM | 128 | 178.258 | 27.567 | 3.745 | 12.813 |
| slow OU | 128 | 178.258 | 35.106 | 13.792 | 50.429 |
| medium OU | 128 | 178.258 | 44.221 | 36.794 | 82.256 |
| fast OU | 128 | 178.258 | 67.237 | 78.888 | 113.169 |
| balanced harmonic increase |  |  |  |  |  |
| BM | 128 | 683.621 | 39.878 | 14.424 | 73.063 |
| slow OU | 128 | 683.621 | 101.417 | 81.108 | 124.902 |
| medium OU | 128 | 683.621 | 127.303 | 123.606 | 127.94 |
| fast OU | 128 | 683.621 | 128 | 127.939 | 128 |
| balanced geometric decrease |  |  |  |  |  |
| BM | 128 | 54.926 | 24.599 | 2.977 | 2.796 |
| slow OU | 128 | 54.926 | 26.783 | 5.735 | 7.878 |
| medium OU | 128 | 54.926 | 28.502 | 10.116 | 14.189 |
| fast OU | 128 | 54.926 | 31.144 | 19.056 | 26.283 |
| balanced geometric increase |  |  |  |  |  |
| BM | 128 | 658.375 | 48.870 | 36.286 | 93.302 |
| slow OU | 128 | 658.375 | 106.934 | 95.213 | 125.693 |
| medium OU | 128 | 658.375 | 127.340 | 123.876 | 127.943 |
| fast OU | 128 | 658.375 | 128 | 127.939 | 128 |
| balanced long root branch |  |  |  |  |  |
| BM | 128 | 7.259 | 24.599 | 2.977 | 1.81 |
| slow OU | 128 | 7.637 | 24.768 | 3.072 | 2.015 |
| medium OU | 128 | 7.637 | 24.902 | 3.175 | 2.233 |
| fast OU | 128 | 7.637 | 25.171 | 3.404 | 2.712 |

**Table 2:** Comparison of relative phylogenetic diversity with the proposed relative pESSs. The values are means from the same 1000 simulations from Tab. 1.

| Model | $n$ | $E[PD/n]$ | $E[n_e^{MI}/n]$ | $E[n_e^E/n]$ | $E[n_e^R/n]$ |
| --- | --- | --- | --- | --- | --- |
| Yule |  |  |  |  |  |
| BM | 15 | 1.007 | 0.475 | 0.263 | 0.367 |
| slow OU | 15 | 1.007 | 0.516 | 0.311 | 0.459 |
| medium OU | 15 | 1.007 | 0.56 | 0.371 | 0.581 |
| fast OU | 15 | 1.007 | 0.641 | 0.5 | 0.675 |
| unbalanced |  |  |  |  |  |
| BM | 15 | 1.031 | 0.556 | 0.303 | 0.615 |
| slow OU | 15 | 1.031 | 0.603 | 0.347 | 0.681 |
| medium OU | 15 | 1.031 | 0.654 | 0.4 | 0.741 |
| fast OU | 15 | 1.031 | 0.753 | 0.526 | 0.834 |
| balanced |  |  |  |  |  |
| BM | 16 | 0.537 | 0.421 | 0.177 | 0.175 |
| slow OU | 16 | 0.537 | 0.447 | 0.198 | 0.225 |
| medium OU | 16 | 0.537 | 0.475 | 0.226 | 0.283 |
| fast OU | 16 | 0.537 | 0.535 | 0.294 | 0.409 |
| balanced short terminal |  |  |  |  |  |
| BM | 16 | 0.677 | 0.368 | 0.168 | 0.081 |
| slow OU | 16 | 0.677 | 0.388 | 0.197 | 0.096 |
| medium OU | 16 | 0.677 | 0.406 | 0.235 | 0.116 |
| fast OU | 16 | 0.677 | 0.438 | 0.317 | 0.161 |
| balanced long terminal |  |  |  |  |  |
| BM | 16 | 2.751 | 0.999 | 0.952 | 1 |
| slow OU | 16 | 2.751 | 1 | 0.977 | 1 |
| medium OU | 16 | 2.751 | 1 | 0.991 | 1 |
| fast OU | 16 | 2.751 | 1 | 0.999 | 1 |
| balanced harmonic decrease |  |  |  |  |  |
| BM | 16 | 1.244 | 0.456 | 0.191 | 0.256 |
| slow OU | 16 | 1.244 | 0.536 | 0.258 | 0.427 |
| medium OU | 16 | 1.244 | 0.632 | 0.356 | 0.602 |
| fast OU | 16 | 1.244 | 0.806 | 0.572 | 0.833 |
| balanced harmonic increase |  |  |  |  |  |
| BM | 16 | 2.481 | 0.703 | 0.391 | 0.7 |
| slow OU | 16 | 2.481 | 0.87 | 0.574 | 0.901 |
| medium OU | 16 | 2.481 | 0.966 | 0.77 | 0.976 |
| fast OU | 16 | 2.481 | 0.999 | 0.961 | 0.999 |
| balanced geometric decrease |  |  |  |  |  |
| BM | 16 | 0.948 | 0.421 | 0.177 | 0.177 |
| slow OU | 16 | 0.948 | 0.471 | 0.222 | 0.276 |
| medium OU | 16 | 0.948 | 0.528 | 0.284 | 0.393 |
| fast OU | 16 | 0.948 | 0.64 | 0.427 | 0.603 |
| balanced geometric increase |  |  |  |  |  |
| BM | 16 | 2.416 | 0.754 | 0.469 | 0.779 |
| slow OU | 16 | 2.416 | 0.896 | 0.637 | 0.922 |
| medium OU | 16 | 2.416 | 0.972 | 0.802 | 0.98 |
| fast OU | 16 | 2.416 | 0.999 | 0.964 | 0.999 |
| balanced long root branch |  |  |  |  |  |
| BM | 16 | 0.225 | 0.44 | 0.177 | 0.12 |
| slow OU | 16 | 0.225 | 0.445 | 0.179 | 0.127 |
| medium OU | 16 | 0.225 | 0.449 | 0.182 | 0.134 |
| fast OU | 16 | 0.225 | 0.459 | 0.188 | 0.149 |
| Yule |  |  |  |  |  |
| BM | 125 | 0.999 | 0.222 | 0.053 | 0.175 |
| slow OU | 125 | 0.999 | 0.246 | 0.098 | 0.324 |
| medium OU | 125 | 0.999 | 0.271 | 0.183 | 0.464 |
| fast OU | 125 | 0.999 | 0.314 | 0.415 | 0.642 |
| unbalanced |  |  |  |  |  |
| BM | 125 | 1.953 | 0.256 | 0.095 | 0.534 |
| slow OU | 125 | 1.953 | 0.292 | 0.151 | 0.683 |
| medium OU | 125 | 1.953 | 0.335 | 0.248 | 0.79 |
| fast OU | 125 | 1.953 | 0.42 | 0.512 | 0.892 |
| balanced |  |  |  |  |  |
| BM | 128 | 0.22 | 0.192 | 0.023 | 0.023 |
| slow OU | 128 | 0.22 | 0.201 | 0.032 | 0.039 |
| medium OU | 128 | 0.22 | 0.209 | 0.045 | 0.061 |
| fast OU | 128 | 0.22 | 0.223 | 0.08 | 0.112 |
| balanced short terminal |  |  |  |  |  |
| BM | 128 | 0.285 | 0.194 | 0.023 | 0.025 |
| slow OU | 128 | 0.285 | 0.205 | 0.035 | 0.053 |
| medium OU | 128 | 0.285 | 0.216 | 0.054 | 0.09 |
| fast OU | 128 | 0.285 | 0.233 | 0.1 | 0.166 |
| balanced long terminal |  |  |  |  |  |
| BM | 128 | 4.806 | 0.973 | 0.703 | 0.998 |
| slow OU | 128 | 4.806 | 0.998 | 0.909 | 1 |
| medium OU | 128 | 4.806 | 1 | 0.984 | 1 |
| fast OU | 128 | 4.806 | 1 | 1 | 1 |
| balanced harmonic decrease |  |  |  |  |  |
| BM | 128 | 1.393 | 0.215 | 0.029 | 0.1 |
| slow OU | 128 | 1.393 | 0.274 | 0.108 | 0.394 |
| medium OU | 128 | 1.393 | 0.345 | 0.287 | 0.643 |
| fast OU | 128 | 1.393 | 0.525 | 0.616 | 0.884 |
| balanced harmonic increase |  |  |  |  |  |
| BM | 128 | 5.341 | 0.312 | 0.113 | 0.571 |
| slow OU | 128 | 5.341 | 0.792 | 0.634 | 0.976 |
| medium OU | 128 | 5.341 | 0.995 | 0.966 | 1 |
| fast OU | 128 | 5.341 | 128 | 1 | 1 |
| balanced geometric decrease |  |  |  |  |  |
| BM | 128 | 0.429 | 0.192 | 0.023 | 0.022 |
| slow OU | 128 | 0.429 | 0.209 | 0.045 | 0.062 |
| medium OU | 128 | 0.429 | 0.223 | 0.079 | 0.111 |
| fast OU | 128 | 0.429 | 0.243 | 0.149 | 0.205 |
| balanced geometric increase |  |  |  |  |  |
| BM | 128 | 5.144 | 0.382 | 0.283 | 0.729 |
| slow OU | 128 | 5.144 | 0.835 | 0.744 | 0.982 |
| medium OU | 128 | 5.144 | 0.995 | 0.968 | 1 |
| fast OU | 128 | 5.144 | 1 | 1 | 1 |
| balanced long root branch |  |  |  |  |  |
| BM | 128 | 0.057 | 0.192 | 0.023 | 0.014 |
| slow OU | 128 | 0.06 | 0.194 | 0.024 | 0.016 |
| medium OU | 128 | 0.06 | 0.195 | 0.025 | 0.017 |
| fast OU | 128 | 0.06 | 0.197 | 0.027 | 0.021 |

**Table 3:** Results of analysis on real data with different definitions of pESS. In the situation where the OU model with disruptive selection (*α* < 0) the value of a was tiny, about 10^−9^. Hence these dynamics on the scale of the phylogeny are indistinguishable from a BM.

| Data set | $n$ | PD/$n$ | $n_e^{MI}/n$ | $n_e^E/n$ | $n_e^R/n$ |
| --- | --- | --- | --- | --- | --- |
| Mantellidae male snout–vent length <sup>1</sup> | 40 BM<sup>2</sup> | 1.364 | 0.411 BM | 0.097 nOU<sup>3</sup> | 0.524 BM |
| Mantellidae Range <sup>1</sup> | 40 OU <sup>4</sup> | 1.364 | 1 OU | 1 OU | 1 OU |
| *Chaerophyllum* fruit length <sup>5</sup> | 33 OU | 0.419 | 0.634 OU | 0.097 nOU | 0.839 OU |
| Carnivora body size <sup>6</sup> | 70 BM | 10.023 | 0.248 BM | 0.052 OU | 0.141 BM |
| Carnivora log(body size) <sup>6</sup> | 70 OU | 10.023 | 0.304 OU | 0.052 nOU | 0.403 OU |
| Carnivora range <sup>6</sup> | 70 OU | 10.023 | 0.498 OU | 0.052 nOU | 0.825 OU |
| Mammalian body mass <sup>7</sup> | 49 BM | 18.48 | 0.125 BM | 0.288 BM | 0.193 BM |
| Mammalian running speed <sup>7</sup> | 49 OU | 18.48 | 0.238 OU | 0.351 OU | 0.432 OU |
| Carnivores, Herbivores hind limb length <sup>7</sup> | 49 OU | 18.48 | 0.319 OU | 0.394 OU | 0.555 OU |
| Carnivores log(body size) <sup>8</sup> | 16 BM | 1.104 | 0.712 BM | 0.362 BM | 0.767 BM |
| Salamanders log(body size) <sup>8</sup> | 197 OU | 0.869 | 0.227 OU | 0.059 OU | 0.375 OU |

<sup>1</sup> [Pabijan et al., 2012] <sup>2</sup> Brownian motion <sup>3</sup> Ornstein–Uhlenbeck with $\alpha < 0$ <sup>4</sup> Ornstein–Uhlenbeck with $\alpha > 0$ <sup>5</sup> [Piwczyński et al., 2015] <sup>6</sup> [Diniz-Filho and Tôrres, 2002, Pavoine et al., 2005a] <sup>7</sup> [Dray and Durfor, 2007, Garland, T., Jr. et al., 1993, Garland and Janis, 1993] <sup>8</sup> [datasets in GEIGER R package Harmon et al., 2008]

Table 3: Results of analysis on real data with different definitions of pESS. In the situation where the OU model with disruptive selection ( $\alpha < 0$ ) the value of $\alpha$ was tiny, about $10^{-9}$ . Hence these dynamics on the scale of the phylogeny are indistinguishable from a BM.
|  |  |  |  |  |  |
| --- | --- | --- | --- | --- | --- |
| Turtles log(body size) <sup>8</sup> | 226 OU | 0.544 | 0.228 OU | 0.233 OU | 0.555 OU |
| Primates log(body size) <sup>8</sup> | 233 BM | 0.513 | 0.181 BM | 0.025 BM | 0.085 BM |
| Darwin's finches log(wing length) <sup>8</sup> | 13 BM | 0.703 | 0.922 OU | 0.292 nOU | 0.941 OU |
| Darwin's finches log(tarsus length) <sup>8</sup> | 13 BM | 0.703 | 0.504 BM | 0.292 nOU | 0.453 BM |
| Darwin's finches log(culmen length) <sup>8</sup> | 13 OU | 0.703 | 0.952 OU | 0.293 nOU | 0.963 OU |
| Darwin's finches log(beak diameter) <sup>8</sup> | 13 BM | 0.703 | 0.504 BM | 0.292 nOU | 0.453 BM |
| Darwin's finches log(gonys width) <sup>8</sup> | 13 BM | 0.703 | 0.753 OU | 0.292 nOU | 0.799 OU |
| Ducks log(brightness) <sup>9</sup> | 38 OU | 0.863 | 0.461 OU | 0.592 OU | 0.684 OU |
| Ducks log(hue) <sup>9</sup> | 38 OU | 0.863 | 0.492 OU | 0.639 OU | 0.724 OU |
| Ducks log(spacing) <sup>9</sup> | 38 OU | 0.863 | 0.692 OU | 0.84 OU | 0.856 OU |
| *Anolis* log(SSD) <sup>10</sup> | 23 BM | 0.97 | 0.61 BM | 1 OU | 0.719 BM |

Currently Ornstein–Uhlenbeck (OU) process are the state of the art in modelling trait evolution [Bartoszek et al., 2012, Beaulieu et al., 2012, Clavel et al., 2015, Cressler et al., 2015, Ingram and Mahler, 2013, Uyeda and Harmon, 2014]. This OU process on a phylogeny is multivariate normal. Therefore all the information will be contained in the mean vector and covariance matrix. In fact we have a natural multiple regression approach and each species, *y*_*i*_, can be represented as

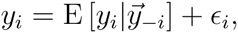

where 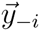 is the vector of measurements without the *i*–th entry. The above equation will be of course of the form

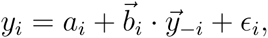

where *ϵ*_*i*_ will be independent of 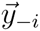. The residual *ϵ*_*i*_ is mean 0, normally distributed with variance

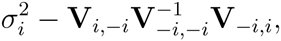

where –*i* notation again means removing the appropriate rows and/or columns. As the variance of *y*_*i*_ is 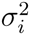, then the independent of the other species part of this variance equals 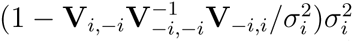. Standardizing every species to variance 1 will mean that each species carries 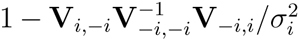 signal specific to itself. Therefore I propose to define a phylogenetic effective sample size, called *regression effective sample size* (rESS), in the following way. Let

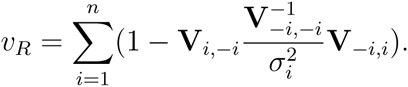

be the total independent signal. The sum *v*_*R*_ can be can be easily lesser than one. We therefore consider

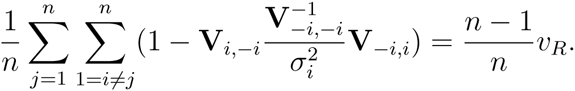

In the above one averages over all species, for each one considering the amount of distinct signal from it. As we know that there is at least 1 species I now define the rESS as

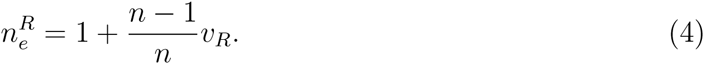

It can be easily checked that 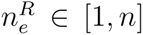, equalling *n* when the species are independent. Taking the pseudoinverse instead of the inverse gives the value of 1 when all *n* species are identical.

The rESS, just as the mESS, can be calculated for any process evolving on a phylogenetic tree. However, just as the mESS does not catch everything about a normal process, the rESS will not catch everything in the non–normal process situation. In the non-normal process, e.g. heavy tailed distributions [Elliot and Mooers, 2014], situation it is necessary to reach for more complicated mathematical tools. The motivation behind the multiple regression approach is to measure how much signal is contained about each species in other species and how much is specific to that species. Another way of formulating the problem is to ask: how much information is contained in the joint distribution of all of the species, when compared with only the marginal distributions. The natural mathematical framework for this is information theory and the concept of mutual information.

As the name itself suggests mutual information quantifies how much information do different probabilistic objects contain about each other. I will briefly introduce a few concepts from information theory pointing the reader to e.g. Koch [2014] for a more detailed discussion.

### Definition 1.

[Koch, 2014] *Let 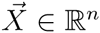 be a random vector with density f such that it has mean 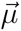 and covariance* V. *Further let f_j_ (j = 1,…, n) be the marginal densities of f and f_G_ be a Gaussian density with the same mean 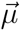 and covariance* **V**, *i.e. for x* ∈ ℝ^*n*^

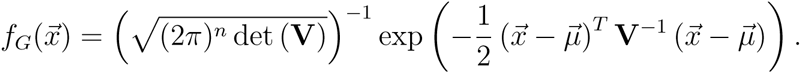

*We then define the following*.

1. *The entropy of f as*

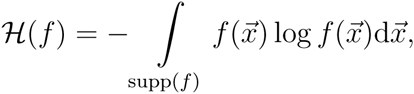

*where* supp 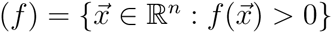 *is the support of f*.
2. *The negentropy of f as*

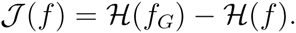
3. *The mutual information of f as*

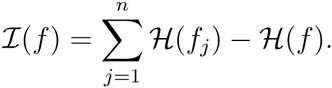

Intuitively the entropy of a density (or rather random variable behaving according to its law) is the measure of uncertainty about the value of this random variable prior to observation. The negentropy from our perspective is more of a technical term however, the mutual information between two densities (or random variables) will be very important in proposing an effective sample size definition.

The maximum sample size attained is *n*, when all species are independent of each other (we have a star phylogeny). In this situation the density function of our *n* dimensional vector of observations will be the product of the marginal *n* densities. No observation contains relatively more information about any other one observation than any other does. Therefore, to quantify how much information do sample points contain about each other, we will consider in Lemma 1 the mutual information between the sample’s *n*-dimensional density and the density defined as the product of the marginal densities. If we recall that all the considered evolutionary models here (Brownian motion, Ornstein–Uhlenbeck) are multivariate normal, then we should expect that the entropy based measures be dependent only on the covariance matrix and marginal variances. In the Gaussian case, all shared knowledge is coded in the covariance structure, see Lemma 1.

### Lemma 1.

[Koch, 2014] Using the notation of Definition 1 the entropy, negentropy and mutual information posses the below properties and relationships between them.

1. The negentropy ***ℐ*** ≥ 0 and ***ℐ***(*f*) = 0 iff *f* is Gaussian.
2. The mutual information ***I*** ≥ 0 and ***I*** = 0 iff 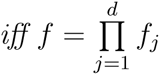.
3. If f is Gaussian, then it has entropy

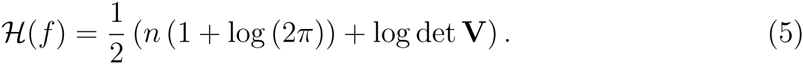
4. If **V** is invertible, then

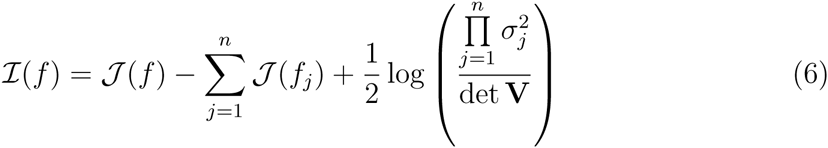

where 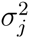 are the diagonal elements of **V** — the marginal variances. If *f* is Gaussian this simplifies to

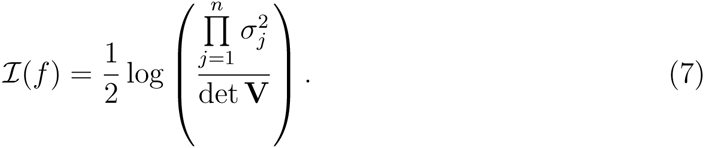 It would be tempting to propose mutual information effective sample size as something

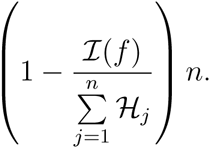

However, ***H**_j_* can easily be negative. We therefore have to find some other way of using the entropy. Lin et al. [2007] used a similar motivation to define an effective sample size in order to obtain correct standard errors for parameter estimates. Theirs was a Bayesian setting and they define the effective sample size as a minimizer of a relative entropy. The relative entropy is between the posterior parameter distribution under the true model and the the posterior parameter distribution under the effective sample. However, their approach does not allow for fractional sample sizes and could require, in the phylogenetic case, optimizing over the power set of species. Therefore, I propose to define the mutual information ESS (miESS) as

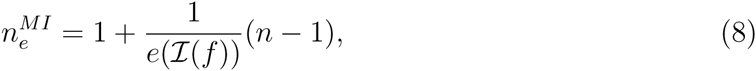

where *e*(·) is a strictly increasing function such that *e*(0) = 1 and *e*(∞) = ∞. One example of such a function is the logarithm of ***I***(*f*) increased by exp(1), considered in this work. I choose such a function as compared to other formulae, e.g. exp(·), it resulted in phylogenetic ESSs similar to those defined by the two other formulae. However, the proposed formula for *e*(·) should only be treated as a temporary definition. Further work is needed to appropriately define it so that e.g. in the case of normal processes (like BM or OU ones) it agrees with the rESS. In order to calculate miESS one needs knowledge of the joint distribution of the tip species, or at least posses a numerical procedure for obtaining it. Both could be unfortunately difficult to obtain in the non–normal case, but [Elliot and Mooers, 2014] present a family of heavy–tailed stable distributions for which the joint likelihood is calculable.

The ESS, defined as such, has the desirable properties of being between 1 and *n*. In the Gaussian the formula for the miESS will equal

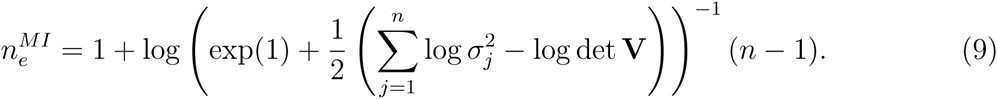

It is important to notice, that the three proposed concepts of effective sample sizes are not compatible with each other. Firstly the mESS is meant to quantify only information about the expected value of the sample, not about independent signal. The motivations behind miESS and rESS are the same, but it remains for a further study to define an appropriate transformation *e*(·) that will make miESS equal to rESS in the normal process case. In Sections 3, 4, 5 and 6 I study their behaviour for simulated and real data.

### 2.1 Multivariate extension

All of the above three definitions assumed that the each of the sample points is univariate. However, methods for studying multiple co-evolving traits on the phylogeny are being developed [see e.g. Bartoszek et al., 2012, Beaulieu et al., 2012, Clavel et al., 2015, Hansen et al., 2008] and all three considered ESS concepts are immediately generalizable to higher dimensions. Assume now that we have a *d* dimensional trait. Each of our *n* points a *d* dimensional observation, our sample is of size *d* · *n* correlated points instead of *n* and **V** ∈ ℝ^*nd*×*nd*^ instead of ℝ^*n*×*n*^. Hence, for model selection purposes we can use the above described procedures replacing *n* with *d*·*n* inside all formulae, as most software packages do.

The miESS and rESS can be elegantly generalized to quantify how many *d*–dimensional observations we have effectively, i.e. how many effectively independent species do we have amongst our *n* species, regardless of the dimensionality of each species. Notice that Eq. (8) does not depend on the dimensionality of the species and can be used nearly without change

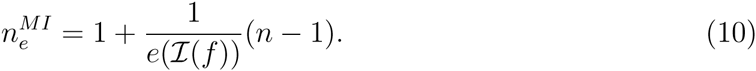

The only difference is that here ***H***_*j*_ is the entropy not of a univariate random variable, but of the *d*_*j*_–dimensional random vector of the *j*–th species. In the Gaussian case, we obtain

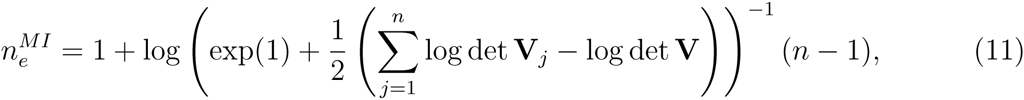

where V_*j*_ is the *j*–th *d*_*j*_–dimensional diagonal block of **V**, i.e. the marginal covariance matrix of the *j*–th *d*_*j*_–dimensional observation.

In a similar fashion we can adapt the 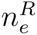 to count the number of effective species in the multitrait case. We sum the conditional total variances i.e.,

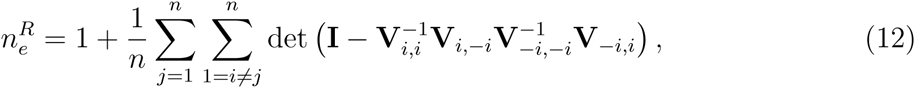

where **I** is the unit matrix of dimension equalling the number of traits. Here –*i* notation means removing rows/columns corresponding to the ith species. Notice that in no case is it required that all species are of the same dimension, allowing for proper handling of missing data.

## 3 Phylogenetic effective sample size

Effective sample size calculation is very important in the phylogenetic context but it seems to have received little attention. Phylogenetic comparative methods have taken care of the inflated sample size phenomena for the most important inference issues. We obtain the correct likelihood value and may in principle obtain correct confidence intervals, and p–values. However, further development is needed for problems that actually depend on the sample size. Effective (and not observed) sample sizes are important when quantifying the biodiversity of a clade to e.g. develop conservation strategies or when doing model selection.

It would seem desirable, to be able to calculate the effective sample size directly from the phylogeny and base any further estimates on this value of *n*_*e*_. In fact, this seems to be the postulated approach by Nunn [Ch. 11 2011], that one should use the tree’s phylogenetic diversity to obtain the amount of information in the sample. Nunn [2011] does not formulate it exactly in this way but this is how mathematically it should be understood. In Section 5, on phylogenetic diversity and conservation I discuss this in detail. However, using phylogenetic diversity to obtain an effective sample size for a trait (or suite of them) will be akin to assuming a Brownian motion (neutral drift) model of evolution. Phylogenetic diversity is the sum of all branch lengths on a tree and this is proportional to the sum of the variances of independent changes on the tree.

However, as Hansen and Orzack [2005] pointed out Brownian change is not appropriate for traits under stabilizing selection. **I** discussed earlier, that all considered definitions of effective sample size will depend on **V**, the between-species covariance matrix, and how it differs from a diagonal matrix. Therefore, we need to calculate *n*_*e*_ based on **V** and not just the phylogeny. The between species covariance matrix depends not only on the phylogeny, but also on the model of evolution. We denote by **T** = [*t*_*ij*_]_1 ≥ *i*, *j*, ≥ *n*_ the matrix of speciation times, where *t*_*ij*_ is the speciation time of species *i* and *j* and *t*_*i*_ the time of species *i* (these will be all equal to the tree height if the tree is ultrametric). Bartoszek et al. [2012] report the form of **V** for various models of evolution.

- Unconstrained evolutionary model — univariate Brownian motion defined by the stochastic differential equation (SDE): d*X*_*t*_ = *σ*d*B*_*t*_

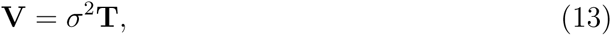 where *B*_*t*_ is the standard Wiener process.
- Constrained evolutionary model — univariate Ornstein–Uhlenbeck process defined by the SDE d*X*_*t*_ = –*α*(*X*_*t*_ – *θ*_*t*_)d*t* ₊ *σ*d*B*_*t*_:

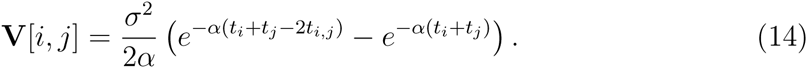
- Multitrait unconstrained evolutionary model — multivariate Brownian motion defined by the 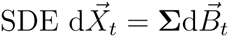:

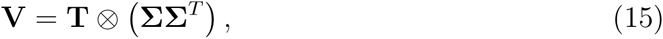

where ⊗ is the Kronecker product.
- Multitrait constrained evolutionary model, traits adapting to constrained traits — multivariate Ornstein–Uhlenbeck process defined by the 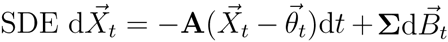

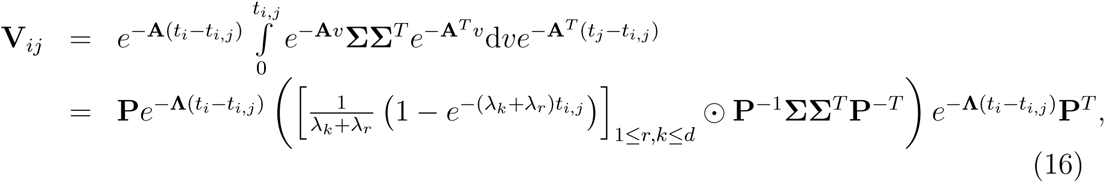

where ⊙ is the Hadamard product, **P**, **Λ** = diag(*λ*_1_,…, *λ*_*d*_) are the eigenvectors and eigenvalues of **A** and **V**_*ij*_ is the block *i*, *j* of dimension *d* × *d* of **V**, i.e. the intersection of the rows ((*i* – 1)*d*,…, *id*) and columns ((*j* – 1)*d*,…, *jd*).
- Multitrait constrained evolutionary model, traits adapting to unconstrained traits — multivariate Ornstein–Uhlenbeck process defined by the SDE system

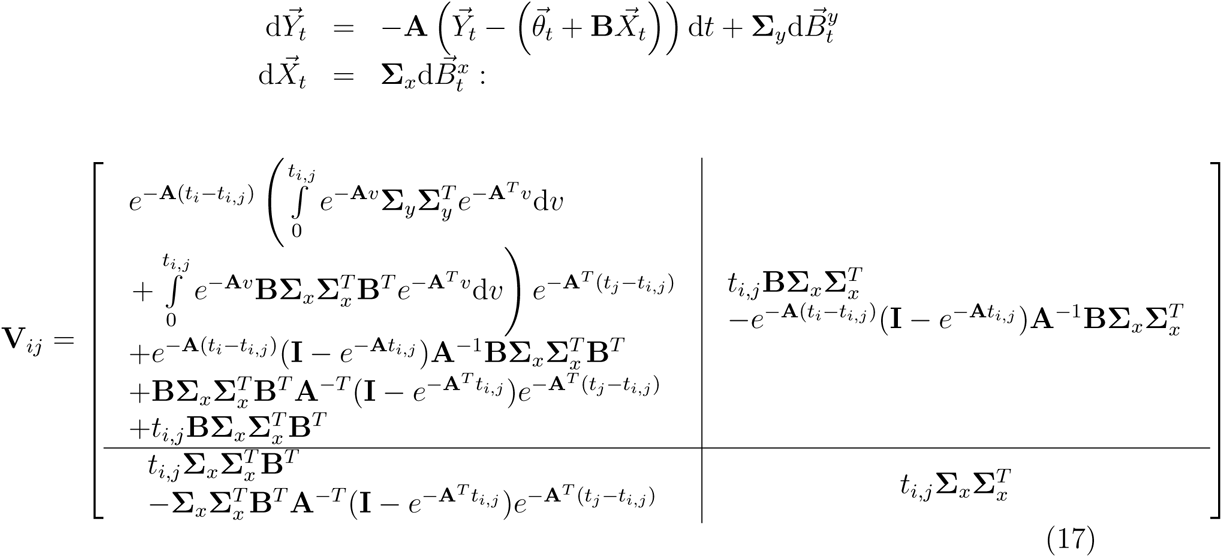

where **I** is the identity matrix of dimensions *d* × *d*.

Hence, before reporting an effective sample size for a clade one has to estimate the parameters of the evolutionary model. It would be also interesting to consider more complex Gaussian setups, like function-valued traits. Jones and Moriarty [2013] consider such a setup: for each species they observe measurements at a vector of coordinates. As they assume normality, jointly the data is multivariate normal, indicating the usefulness of all three proposed pESSs.

Given a phylogenetic tree and model of evolution, we can easily calculate the effective sample size by plugging in the appropriate formula. Below I present the values of the different definitions of ESS for the BM model of evolution. Formulae for OU based models would be too lengthy to be readable. We assume the tree is ultrametric with height *T*.

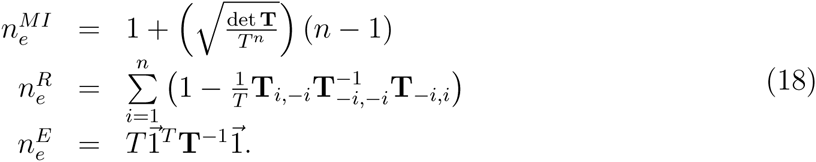

In the phylogenetic context it would be tempting to take for the ESS factor, *p*, the interspecies correlation coefficient [Sagitov and Bartoszek, 2012]

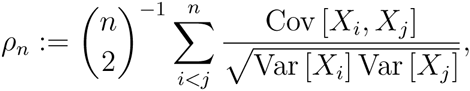

where the sum is over all pairs of tip species. The above random variable is very well studied for the pure birth tree. The expectation of *ρ*_*n*_ was derived for the BM and OU process [also with jumps Bartoszek, 2014, Bartoszek and Sagitov, 2015b, Sagitov and Bartoszek, 2012]. Recently Mulder and Crawford [2015] calculated the distribution under the above modes of evolution. However, in all the considered models E[*ρ*_*n*_] → 0. Furthermore, for BM on a tree with extinction one can consider death coefficients such that E [*ρ*_*n*_] → 1. As 0 ≤ *ρ*_*n*_ ≤ 1, by the dominated convergence theorem, we have *ρ*_*n*_ → 0 (alternatively → 1) almost surely. Such almost sure 0 or 1 asymptotic behaviour is not consistent with the motivation behind studying a pESS, where the sample should be somewhere between 1 and *n*, not exactly 1 or *n*.

I illustrate Eq. (18) in Fig. 2. I also include the effective sample sizes for Ornstein–Uhlenbeck models. The considered evolutionary scenarios are a Brownian motion and Ornstein–Uhlenbeck process. We fix the initial state *X*_0_ = 0 and *σ*^2^ = 1. For the OU process we also fix the optimum *θ* = 1. We vary the adaptation rate *α* = 0, 0.25, 0.5,1. We consider three binary phylogenetic tree setups (see Fig. 1). Two are deterministic trees: a completely unbalanced tree, a completely balanced tree (number of tips is a power of two). The third type is a random one — a conditioned on the number of tip species Yule (pure birth) tree [Bartoszek and Sagitov, 2015b, Gernhard, 2008a, b, Sagitov and Bartoszek, 2012]. The rate of speciation is taken at *λ* = 1. I take the number of tip species to be from 5 to 200. Of course in the balanced tree only those that are powers of two are allowed, hence there were significantly fewer trees. Each point is the average over 1000 simulations.

To make the simulations comparable the heights of the two deterministic tree types were scaled to log n, the expected height of the Yule tree. Also for these topologies randomness was added by drawing the length of the root branch from the exponential with rate 1 distribution. In the case of the OU model, it allows the process to approach stationarity/stasis before speciation starts to take effect.

## 4 Phylogenetic information criteria

My main motivation for studying the effective sample size in the phylogenetic context is obtaining correct values of information criteria that depend on sample size. Information criteria are necessary for e.g. finding the best evolutionary model, testing evolutionary hypotheses, distinguishing between competing phylogenies [Bartoszek and Lió, 2014] or regime layouts [Butler and King, 2004]. If the evolutionary models/hypotheses are nested, then models can be compared by a likelihood-ratio test. Such a test tells us whether the increase in the number of parameters significantly improves the model fit. Alternatively when the models are not nested the Akaike information criterion that penalizes for the number of extra parameters

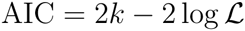

was proposed [Akaike, 1974]. In the above *k* is the number of parameters and ***L*** the likelihood. The model with the lower AIC value is the better one. However, both the *χ*^2^ distribution of the likelihood ratio test and the AIC are asymptotic approaches. They will be correct when the sample size is infinite (or large in practice). In phylogenetic comparative studies the number of species is usually small. Therefore two alternative criteria that correct for small sample size were proposed to the phylogenetic comparative methods community [Hansen et al., 2008]

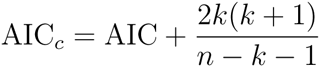

and the Bayesian (or Schwarz) information criterion [Butler and King, 2004]

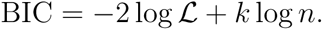

Of these two the AIC_*c*_ seems to be the more used one (but AIC is also very popular).

To see how much of a difference it makes, whether the observed or effective number of species is used, I performed a simulation study under various evolutionary scenarios. Under each scenario I simulate data *N* = 1000 times and from this obtain histograms of the AIC_*c*_ values under the true model and an alternative using both the number of species and the effective sample size, Figs. S.1—S.8 in the supplementary material. I also plot in Fig. 3 how the average value of the small sample size correction changes under the different evolutionary models and effective sample size value. We consider the same evolutionary scenarios as in Fig. 2 and observe that for large *α* identifiability of the true model is easier. The histograms of the AIC_*c*_ are shown for small (*n* = 30) and large (*n* = 205) phylogenies. We can see that in the large phylogeny case, all definitions of sample size result in the same distribution of AIC_*c*_. However for the small phylogeny the mean and regression ESSs, 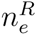 and 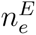, seem to be more effective with the balanced phylogeny and fast adaptation. The simulation results furthermore show that distinguishing different adapting OU models from each other and the BM one can be difficult. This difficulty, especially with smaller αs, is to be expected as the slowly adapting processes can take a lot of time to reach stationarity and loose ancestral signal [Adamczak and Miłoś, 2014, 2015, Ané et al., 2014, Bartoszek and Sagitov, 2015b]. In fact our simulations confirm in this respect Cressler et al. [2015]’s recent study — “Selection opportunity (*i.e. α*) is substantially more difficult to estimate accurately: … relative errors exceeding 100% are common, even when the correct model has been selected.” [especially for small *n* and *α*, see Fig. 6 of Cressler et al., 2015]. Hence, significantly larger sample sizes would be needed to identify slowly adapting models. Figure 3 also tells us that even with smaller sized phylogenies all pESS definitions should result in similar AIC_*c*_ values. The observed agreement, between all tested sample size definitions, suggests that the likelihood dominates the AIC_*c*_, which is not surprising as the data is simulated under the BM or OU models. A similar consistency is observed when working with real data (Section 6). The situation is different for the fully balanced tree which holds the most dependencies between the species. In such a symmetric case, probably a much larger tree would be needed to obtain stability.

**Figure 3:**
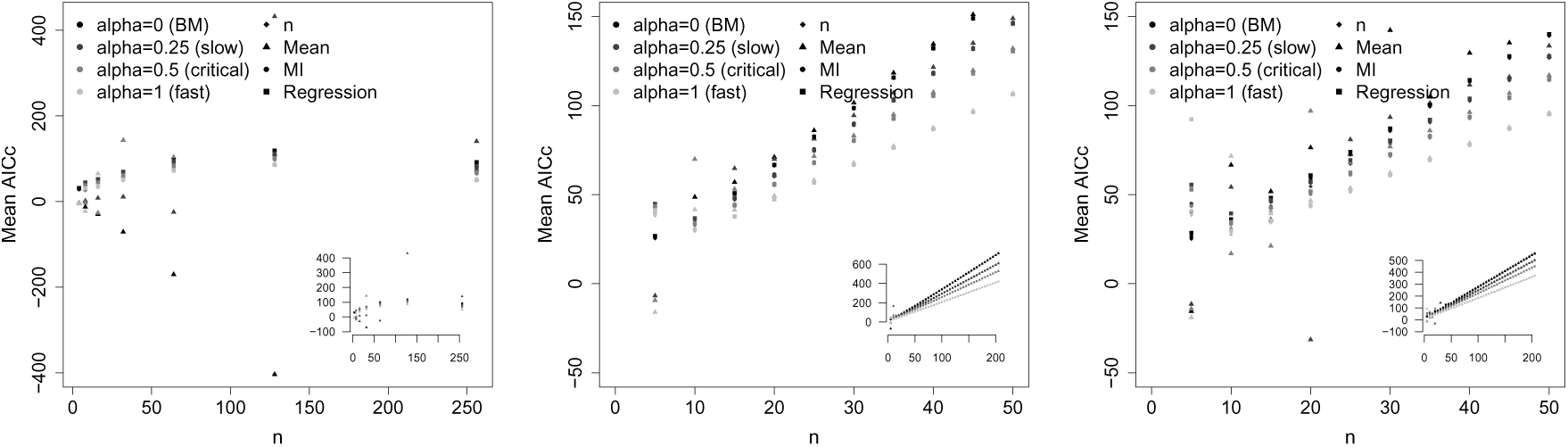
AIC_*c*_ effective sample size correction for different types of trees and evolutionary processes. Left: balanced tree, centre: left unbalanced tree, right: average of 1000 pure–birth Yule trees (*λ* = 1). The balanced trees and unbalanced trees were generated using the function stree() of the R ape package, the Yule trees by the TreeSim R package. The parameters of the processes are Brownian motion (*X*_0_ = 0, *σ*^2^ = 1), second row: Ornstein–Uhlenbeck process (*α* = 0.25, *σ*^2^ = 1, *X*_0_ = 0, *θ* = 0), third row: Ornstein–Uhlenbeck process (*α* = 0.5, *σ*^2^ = 1, *X*_0_ = 0, *θ* = 0), fourth row: Ornstein–Uhlenbeck process (*α* = 1, *σ*^2^ = 1, *X*_0_ = 0, *θ* = 0).

Ané [2008] noticed that for a Brownian motion model of evolution effective sample sizes can be very small. Garland, T., Jr. et al. [1993]’s mammal phylogeny had 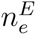 = 6.111 with 49 tip species. My simulations give very similar numbers (Fig. 2). A Yule tree of 50 tips has E [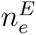] = 5.391, E [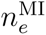] = 14.574 and E [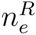] = 11.455, a fully unbalanced tree with 50 tips has E [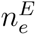] = 7.781, E [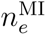] = 17.06 and E [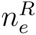] = 27.802 and a fully balanced tree of 64 tips has E [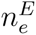] = 2.909, E [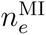] = 9.729 and E [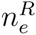] = 2.8.

The very low amount of independent information is evident. In Section 6 I reanalyzed Garland, T., Jr. et al. [1993]’s mammalian data [from the ade4 R package Dray and Durfor, 2007]. Of course 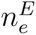 = 6.111, as expected for the mammalian body size evolution (the BM model was selected). The other pESSs were not much higher 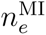 = 14.125 and 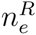 = 9.437 (also BM model). In Section 6 I discuss this data set in more detail.

In most cases, the mean effective sample size is the lowest because it measures the information that the sample contains on the mean value. In the BM case, this is the ancestral state and there is very little information on it. The other pESSs look more holistically at what dependencies are in the data and hence are larger. If we move to more and more adaptive OU models (increase *α*), then all, but especially 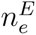, increase. The mean ESS is nearly always the smallest. However, if adaptation is fast and terminal branches are long (i.e. the contemporary sample is nearly independent), then it can also be nearly *n* (see Tab. 1).

Based on the simulation results alone, it is difficult to provide rules of thumb for the applied user. All methods essentially give the same results (as they should under simulated data!). However the analyses of real data in Section 6 does provide some recommendations which are there discussed. One suggestion from the simulations is that it is not that important which information criterion one uses — all should result in the same conclusion. In the PCM field there is a tradition to prefer the AIC_*c*_ and BIC over the AIC, but at least in this study I did not notice significant differences.

## 5 Phenotypic diversity and conservation

An important application of phylogenetic methods is to quantify the biodiversity of a group of species. Phylogenetic methods allow one to formulate definitions of species that are useful from an evolutionary point of view [Ch. 11 Nunn, 2011]. I will not be concerned with a definition of a species but assume that some phylogeny relating predefined taxonomic units is available. The impact of a phylogenetic definition of species was investigated by Agapow et al. [2004] and they noticed that this caused an average increase of the number of species by about 49% when compared to alternative definitions. Such an influx can mean that a lot of species were “split”, resulting in species with smaller populations and geographic ranges. In turn as these are variables contributing to classifying a species as endangered, it may lead to more labelled as such. Therefore Agapow et al. [2004] postulated quantifying conservation value using alternative variables, one of which was trait diversity.

Faith [1992] suggested to quantify biodiversity through phylogenetic information. The main idea is that one should concentrate on feature diversity — how diverse are organisms. Diversity is of course something difficult to quantify, we do not even know of all the variables to measure. Crozier [1997] pointed out that one of the aims of conservation is to “maximize the preserved information of the planet’s biota best in terms of genetic information”. He then points out that phylogenetic based measures which include branch lengths will be better indicators than just counting the number of species. Therefore as a proxy Nunn [2011] following Faith [1992] proposes [but also refers the reader to Faith, 1994, 2002, Crozier, 1997, Purvis et al., 2005] to quantify feature diversity with phylogenetic diversity (PD) — the sum of branch lengths of a tree/clade. The extinction of a clade (or species) is therefore equivalent to subtracting the amount of branch lengths particular to this clade. Phylogenetic diversity has a rich literature [e.g. Crawford and Suchard, 2013, Mooers et al., 2012, Stadler and Steel, 2012] and therefore it is possible to make quantitative predictions about diversity loss/retention under different models of tree growth, extinction and conservation.

From a mathematical perspective PD quantifies the amount of feature diversity as the amount of accumulated variance under the assumption that evolution follows Brownian motion. One may say that this is sensible as an overall feature variable describing a species will be the sum of effects of many traits. Individually traits may be under selection but their sum is not necessarily adapting to anything — providing an argument for Brownian drift.

An alternative approach that could be used to quantify the biodiversity (or feature diversity) of a clade of *n* species is the effective amount of species in this clade *n*_*e*_. This is done in a straightforward way. We prune the phylogeny to the subtree which contains only this clade, and use the methods described in this work to obtain *n*_*e*_ for this subtree. Such an approach could be more appropriate for various reasons. For example it could turn out that the traits important from a conservation point of view are quantified by another process e.g. Ornstein–Uhlenbeck. In the OU case, the changes along disjoint parts of the phylogeny are not independent and the variance is not a linear function of time.

The above trait based approach for quantifying biodiversity is closely related to the ideas presented by Pavoine et al. [2005a]. They introduce the “originality of a species within a set” concept based on Rao [1982]’s quadratic entropy that describes the “average rarity of all the features belonging to this species” [see also Pavoine et al., 2005b]. In the discrete trait case, Pavoine et al. [2005a] find it equivalent to phylogenetic diversity. They analyze the Carnivora data set [Diniz-Filho and Torres, 2002, Pavoine et al., 2005a] and plot (their Fig. 3) how the PD changes with the amount of species dropped. Interestingly the PD reaches a final plateau around 58 (out of 70) species — the same amount that is the rESS for the range measurement.

Phylogenetic diversity measures can be overturned if one uses the diversity of a (suite of) trait(s) as a proxy for biodiversity. In a very wide and recent shallow radiation the diversity of a trait can be very small while the sum of branch lengths can be large. On the other hand if we have trees with few very old tips, then they may have much lower phylogenetic diversity. However, they might have diverged so far back in time and accumulated so much change in their phenotype (without speciating), that the loss of even one tip results in a much more significant loss of phenotypic innovation than even of most of a recent shallow radiation. The latter is intuitively obvious — in a recent shallow radiation the majority of information about all species is coded inside essentially all of the species. It suffices for only one to survive for most of the information to be retained. But making the radiation wider and wider one can imagine increasing the measure of phylogenetic diversity as much as desired. Of course loosing tips is equivalent to loosing small innovations that set the species apart. All changes are naturally a value in themselves but the majority of information is stored in any individual tip. However, in the many old species case, every single species is a distinct entity not containing much information about the rest. Hence, any loss of a single species leads to an irreplaceable loss of diversity [Nee and May, 1997], while the phylogenetic diversity measure might not pick this up. Nunn [2011, p. 319] points out that we are losing biological and cultural diversity at a faster rate than ever before. Therefore, it is important to quantify how much of what we loose.

Rather recently Vellend et al. [2011] compared various phylogenetic based measures of biodiversity, including PD. They found that mean PD (mPD, average over all pairs of species phylogenetic distance) was more sensitive in detecting “non-random community assembly” in a clade. This is probably due to mPD taking advantage of more information, the branch lengths and tree topology (averaging over pairs).

The pESS can be considered as a proposal of a new multi-omics currency of biodiversity. Instead of the standard currencies “species” or PD I use diversity in traits. In other words, I sum up innovations particular to species. Based on such a partition of the variance one can identify “innovative” clades which contain a lot of information. The proposed in this work approach can be a step towards species-free methods postulated by Agapow [2005]. As yet, the pESS is not completely species-free of course, it still includes the phylogeny. The tips of the tree are pre-defined by experts taxonomic units. However, it is not an only-species methodology as e.g. counting species would be. It includes evolutionary process information, that takes into account the topology of the tree — how much of one species is there in another. Also, Agapow [2005] discusses that the problem with species methodologies is that depending on the definition of species we can get wildly different counts. Isaac and Purvis [2004] point out that correct identification of species numbers is important for understanding the diversity of our world.

Therefore, if one misidentifies a species, problems could occur — the species count will be wrong and hence the phylogenetic diversity. It will be based on too few or too many branches. And what if one missed a particular subpopulation that had something very special attached to it? Can one still include its diversity even though it does not appear on the phylogeny? The pESS can precisely do this through integrating data from different sources. Assigning the effective clade size that takes into account the phylogeny and trait variance and between-species covariance, should allow one to strike a balance between expert knowledge concerning species and uncertainty attached to correct demarcation. For example evolutionary models can be easily extended to include intra-species variance, often called “measurement error” [e.g. Felsenstein, 2008, Hansen and Bartoszek, 2012, Rohlfs et al., 2013, to name a few]. Mathematically, these methods boil down to adding to the matrix **V** a matrix **M** which is the intra-species variance (“measurement error”). Then this new covariance matrix **V** ₊ **M** can be treated as the old **V** to obtain a value of effective species number. The intra-species variance can be a representation of our uncertainty about species demarcation and be used to correct for species miscall. If a species has many subpopulations, that are very diverse, representing a species by only its mean over all (measured) individuals will not be the best option. Including the variability of the trait inside the species can partially alleviate the need to know the correct species structure. Such “observational error” can be thought of as averaging over all possible species demarcations that we are not sure about.

Mooers et al. [2005] discuss that one can look at conservation from an ethics point of view should all species be considered equal and protected in the same way or should one protect the features of evolution that are of some value for us. Then phylogenetic diversity is a measure that quantifies a particular feature of evolution. What I propose in my work is quantifying a different feature of evolution. What sets it apart from PD is that it requires the researcher to define traits — exactly what features of evolution are valuable. To illustrate the statement, Nee and May [1997] point out that the loss of *Homo sapiens* would result in a loss of a tiny fraction of evolutionary history, when one uses a measure that takes into account only the tree. If we would choose a trait associated with e.g. civilization achievements and then calculate the ESS of the human lineage (1 by definition) and non–human clades we would obtain a completely different result.

In a way one could say that this is merely replacing counting species with counting the effective number of species. However, the difference is in how we count. Counting just the number of species means enumerating taxonomic units according to some definition. Counting the effective number of species, in the way I propose, is really saying how much biodiversity we have in a clade, where biodiversity is represented by some (suite of) trait(s). This measure can also be thought of as calculating how much innovation we have in the clade. Of course my approach shifts the responsibility to the biologist to identify what traits are important.

Jetz and Freckleton [2015] have very recently published an analysis that is distantly related to what I discuss. They notice that on many species we have too little data, to say if they are endangered or not. On the one hand this would mean that we could assume that all data–deficient species are endangered, but as Agapow [2005] pointed out this would be far too costly. On the other hand Jetz and Freckleton [2015] point out, that Butchart and Bird [2010] observed that data-deficient birds are at no greater extinction risk, than assessed birds. This suggests, that one could use, as Jetz and Freckleton [2015] do, e.g. body-mass, to predict threat status/threat probability. Of course, as species are dependent, in such an analysis the phylogeny needs to be accounted for. Such an approach has the drawback, as Jetz and Freckleton [2015] discuss, that a logistic regression, i.e. threatened/not threatened, will require a large dataset. Therefore, it might be possible, but this of course requires further development and linkeage with phylogeographical models, that effective clade size could also be a proxy for threat status. In addition Jetz and Freckleton [2015] point out, that many species have missing measurements on phenotypes. The evolutionary models used to obtain pESS can handle unobserved data in a natural manner. There is no need to remove a species from an analysis even if it has missing data.

In Tabs. 1 and 2 I present situations where the pESS approaches produce results which are in agreement and disagreement with phylogenetic diversity. I considered a number of different phylogenies (see Fig. 4), with recent shallow radiations, with long tip branches, short tip branches and Yule trees. Two considered types for balanced trees are geometrically or harmonically increasing or decreasing branch lengths. In the geometric case, each level’s branch is half of or twice as (decrease or increase) the previous level’s one. In the harmonic case, the branch length of the *i*–th level (counting from the root — decreasing or from the tips — increasing) is 1/i of the tree’s height. On top of all trees I considered the BM process and the OU process with different parameter values. All trees have an expected height of log *n*. In deterministic trees (balanced and unbalanced, i.e. non–Yule) some randomness to the topology is added by a root branch of length distributed exponentially with rate 1. This is so that the models are more comparable — that some variance is attached to the trait evolution and the OU model is allowed to approach stationarity/stasis before speciation effects begin. For each setup 1000 simulations were made.

**Figure 4:**
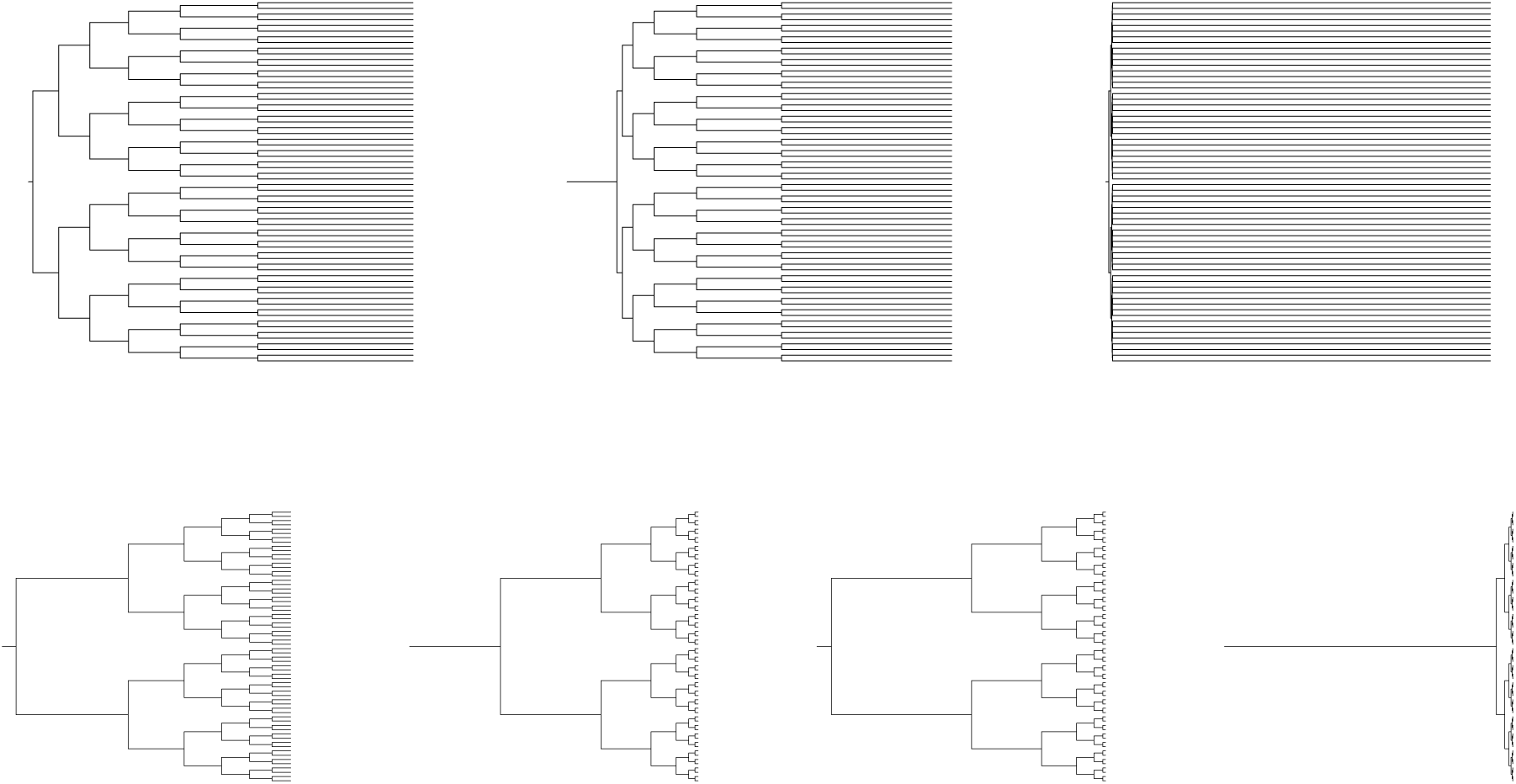
Balanced phylogenies used in pESS for biodiversity simulations. Top row, left to right: branches increasing towards root in a harmonic fashion, branches increasing towards root in a geometric fashion, terminal branches are 99% of tree height. Bottom row, left to right: branches increasing towards root in a harmonic fashion, branches increasing towards root in a geometric fashion, terminal branches are 1% of tree height, root branch is 95% of tree height. The number of tips is 64. For other types of phylogenies see Fig. 1.

The first thing that can strike us in Tabs. 1 and 2 is that PD can be identical despite very different topologies, dependencies and tip species numbers. For example the Yule and unbalanced trees have nearly identical PDs for *n* = 16 while the pESSs suggest that there is a difference between their information content. On the other hand when *n* = 125 there is a large difference between the PDs, while not that much in the pESSs.

If we compare the balanced short terminal tree with *n* = 128 and the *n* = 16 balanced harmonic/geometric increase trees, then they have nearly identical PDs. Their pESSs are also similar but they explain what is going on, in the first case, we have many very similar species in the second a few very distinct ones. In the latter situation, as discussed previously, the loss of a species means loosing a completely separate entity, in the former all species contain significant information about all the others.

Phylogenetic diversity’s lack of explanatory power of the dependency structure induced by the different topologies, is even more evident when considering relative PDs and pESSs, i.e. PD/*n*, *n*_*e*_/*n*, (Tab. 2). In the first example above (unbalanced and Yule) the relative regression ESS seems stable (similar growth with *α*) when comparing the small and large phylogenies (both Yule and unbalanced). It clearly shows that there is more independence in the unbalanced tree — as expected there are more long terminal branches. The relative PD does not distinguish between the small Yule and unbalanced phylogenies, and the large Yule phylogeny, while 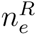/*n* = 0.367, 0.615, 0.175 for small Yule BM, small unbalanced BM and large Yule BM respectively. The regression ESS clearly shows how the tree influences the dependency structure of the tips. Unfortunately the mutual information and mean ones do not describe these dependencies so clearly. However, in their case this is explainable — the mean measures only information on the expected value and the MI one needs further refinement with respect to the *e*(·) transformation. Vellend et al. [2011, p. 208] comment that distance based metrics (e.g. PD) make it easy to detect phylogenetic clustering but not overdispersion on balanced trees, while the opposite is true for unbalanced trees with accelerating diversification. Given an appropriate trait, the pESS should not have such topology dependent problems as uses both phylogenetic and “evolution on a lineage” information. If the species are clustered, then this should be reflected in more dependencies between the observations and lower *n*_*e*_. On the other hand overdispersion should lead to more independence and hence higher *n*_*e*_.

The general pattern from Tabs. 1 and 2 is that if there is a lot of independence, then PD will be large. But as said, the sum of branch lengths does not capture everything. For example, I look in more detail at the balanced long terminal, harmonic and geometric increase topologies. The PD measures (absolute and relative) do not distinguish between these different situations. However, in Fig. 4 we can see that there is a substantial difference between the long terminal one and the harmonic and geometric increases. The long terminal sample should essentially be independent, while the other two should exhibit dependencies. The 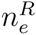 describes such a pattern perfectly. On the long terminal tree all processes generate a nearly independent sample with the rESS measure. For the other two the process has to evolve quickly to loose ancestral dependencies. But on the other hand, by the PD measure the long terminal branch tree carries less independence (diversity) than the harmonic and geometric increase trees. Furthermore it is interesting to notice that the growth of the relative pESSs with α is similar for all pESS definitions. The geometric increase has larger pESSs due to the longer terminal branches.

## 6 pESS in biological data sets

Using the new version of mvSLOUCH I analyzed a number of data sets to see what effects using different definitions of pESS would have on inference. The data sets are a collection from various sources. All but one are vertebrates. The sole exception is the fruit length, a fitness related trait, data for 33 *Chaerophyllum* species [Piwczyński et al., 2015]. Ten datasets from the animal kingdom are looked into. I consider Madagascar Mantellidae male snout-vent length and range measurements for 40 species [Pabijan et al., 2012], Carnivora body size (natural and log scale) and range data for 70 species from the `carni70` data set, ade4 R package [Diniz-Filho and Torres, 2002, Pavoine et al., 2005a]. I look into the same data that Ané [2008] used to introduce what I called the mESS: body mass, running speed and hind limb length of 49 mammalian (both carnivores, herbivores) species [Dray and Durfor, 2007, Garland, T., Jr. et al., 1993, Garland and Janis, 1993]. I also consider the data sets attached to the GEIGER R package [Harmon et al., 2008]: log body size of 16 Carnivores species, log body size of 197 salamanders species, log body size of 226 turtles species, log body size of 233 primates species, and log wing, tarsus, culmen lengths, log beak diameter and log gonys width 13 Darwin’s finches species. I also look into log brightness, hue and spacing of 38 Duck species [Eliason et al., 2014]. Lastly I also use log sexual size dimorphism of 23 *Anolis* species [Butler et al., 2000, Butler and King, 2004].

Data are analyzed on the natural scale unless mentioned above. The results of this analysis are presented in Tab. 3. In all datasets the phylogenetic trees are ultrametric. All trees were rescaled to a height of log(*n*) – 1 to be comparable with other results here. I take the – 1 as there is no root branch in these trees. In all the analysis, except the mammalian hind limb length, the OU processes were assumed to have a single constant optimum over the phylogeny.

As common in many comparative studies BM was selected for the body size/mass variables. There was one exception to this, the logarithm of body size for the Carnivora data gave more support to an OU process. On the other hand measurements on the natural scale are in favour of a BM.

All definitions of pESS, except for the mESS, lead to the same conclusions. Using the mESS can lead, at first sight, to dramatically different conclusions — an Ornstein–Uhlenbeck process with disruptive selection (i.e. *α* < 0). However when looking into the estimate of α in all cases, it was negative but very close to 0 — hence resembling a Brownian motion. Also mESS is not, as explained in the beginning, designed to measure how much independent signal there is in the data. It measures how much information there is in the data to make inference about the mean value parameters. The quantification of the independent signal depends rather on the covariance between the data points — hence the regression and mutual information ESSs seem to make more sense. A reader might ask how is it possible that a more complex model (nOU) is chosen when the mESS is significantly smaller than n. But the mESS for BM models in these situations is even lower hence the disruptive OU model is favoured. However, the α parameter is estimated at the magnitude of –10^9^ so effectively this is a Brownian motion. A similar phenomena can be observed in the *Anolis* SSD analysis. With the mean ESS the more complex OU is chosen as 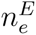 ≈ n. Such a choice is made by the model selection procedure, as under the BM model, 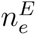 ≈ 5.199.

From these results one can draw the conclusion that even with noisy “real–world” data the likelihood should still be expected to dominate. However, the mESS will not be a fortunate choice to use especially if the data seem to follow a BM. There is a very good explanation for low values of 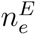. Under the phylogenetic BM model inference about the ancestral state are next to impossible from only the contemporary sample. Due to the noise level one cannot obtain consistent estimators of it [Ané, 2008, Bartoszek and Sagitov, 2015a, Sagitov and Bartoszek, 2012]. As the mESS measures the amount of information available to estimate mean parameters and the ancestral state equals the mean in the BM model, then 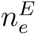 will be small. Hence, AIC_*c*_ will be high in this case, and this model will not be favoured. However, with other definitions of pESS, Brownian motion is not discriminated in this way. When the true model is the OU one, the mESS does not seem to lead to wrong conclusions. This is, as in the OU model there is a lot of information about *θ* — approximately the mean value [Bartoszek and Sagitov, 2015b].

If we look at the turtles and primates results, then we can again see that the PD does not tell the full story of diversity. Both have similar relative (and absolute) PDs but their 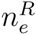 are very different. The primates body size follows a Brownian motion and the phylogeny highly correlates contemporary species. The turtles’ body size on the other hand follows an OU process and there is much more independence in the data set. This is despite the fact that when investigating the phylogenies the primates’ one has clades diverging further back in the past.

An interesting data set to look at are the 49 mammalian (Carnivores, Herbivores) measurements [Garland, T., Jr. et al., 1993, Garland and Janis, 1993] that can be found in the ade4 R package [as the `carni70` data set, Dray and Durfor, 2007]. This was the data that Ané [2008] used to illustrate her work. In line with her conclusions, as mentioned before, I found that the body size variable has a very low amount of independent information. However, the two other considered variables running speed and hind limb length are more informative, with the rESS being more than half *n* for the latter variable. Hind limb length is also interesting that the best supported OU process has different optima for the carnivores and herbivores. On the other hand running speed supported a common optimum for the two groups of mammals.

I ended Section 4 by writing that the analyses of real biological data sets would be better for providing rules of thumb for what how best to use information criteria in a phylogenetic context. Essentially all performed analyses indicate that the choice of criterion and whether to use the observed or effective sample size does not have much effect on model selection. However, unlike in the simulation study, 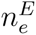 can give very different results. Seeing as this pESS often points to a disruptive OU process, while the other definitions to a more biologically realistic BM or adaptive OU process, indicates that 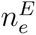 is probably not a good choice for model selection purposes. This seemingly undesirable behaviour, alongside tiny effective sample sizes, of the mESS occurs for samples as large as 70.

A proposed rule of thumb is that if one has a very small sample (like the 13 species for Darwin’s finches), then it is worth trying out the different definitions of ESS for model selection. Of course with such a small sample drawing any conclusions is risky. However, sometimes it is impossible to collect more measurements. The pESS approach might allow the user to look at the observations from different angles in such a data deficient situation. When the sample size is moderately large all methods (bar the mESS based ones) seem to be robust and lead to similar conclusions. Of the three proposed definitions of pESS I found that the rESS performed best. It has furthermore the advantage of a solid mathematical explanation on how it quantifies independence in a phylogenetic data set. However, I only tested it for Gaussian processes. In a non–Gaussian setting the miESS could work better — a topic for further investigation.

## 7 Discussion

In this study I approached the question of quantifying the amount of independent signal in a phylogenetic data set. I proposed two definitions of an effective sample size and compared it to the one considered by Ané [2008]. My work is mainly heuristic — to see how do these proposed definitions behave on real and simulated datasets.

The most important goal of my paper is — does it make sense to use information criteria for model selection with phylogenetically correlated data. The most popular information criterion, Akaike’s, is an asymptotic one with infinite sample size. Because phylogenetic samples are usually small this was not satisfactory — e.g. more realistic but parameter richer models are rejected in favour of simpler ones. Therefore small sample size corrected criteria were implemented, e.g. the considered here AIC_*c*_ (BIC an alternative one). However, these corrections were derived under the assumption of independence. One of the aims of this paper is to propose a formula that allows for replacing the sample size with the amount of independent observations and then see if this changes the models indicated by the criterion. In most cases, it seems that the likelihood part of the information criterion dominates and all definitions of pESS lead to similar conclusions especially with many tip species. One can assume therefore, that for model selection, dependencies in the data do not cause serious problems. However for small phylogenies it seems reasonable to compare the conclusions from different pESS definitions (Tab. 3 Darwin’s finches OU conclusion for 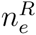 and BM for *n*).

The second goal of the paper is to quantify the amount of dependency in a phylogenetic sample and to understand patterns associated with it. Obtaining the pESS of clades, can indicate clades where more sampling or research effort is needed. For example, is a low pESS due to there being really few species or should we expect more or possibly a reclassification of species is needed? Of course, all of this is with respect to a specific trait(s). This specificity allows for identification of interesting clades. Considering a trait like body size we obtain the distribution of relative (for comparability between clades) pESSs across a set of clades. In the next step one may identify outlier clades — extremely high or low pESSs for further research. Low relative pESSs could indicate recent radiations or other factors not allowing different species to evolve independently. High relative pESS, especially close to 1, would mean that the species are under completely independent evolutionary pressures. Phylogenetic ESSs of a clade can indicate undersampling of species. If we have high relative pESS with a low absolute number of species, then perhaps the very recently evolved species are missing. This can be helpful to indicate where biologists and taxonomists should put efforts to fill in the gaps [Isaac and Purvis, 2004].

A possibly appealing application of this measurement of independence is the quantification of biodiversity. The most commonly used evolutionary measurement tool is phylogenetic diversity — the sum of branch lengths. It seems however that this number does not say much (even when scaled by the number of tip species) about the “value” of an individual species and comparison between clades is difficult (very different ones can have identical values, cf. Tabs. 1 and 2 long terminal with geometric and harmonic increases, or Tab. 3 primates and turtles). Therefore, to give the “value” of a single species I propose to use the relative pESS (i.e. *n*_*e*_/*n*). If the value is low, then the loss of a single species does not result in much biodiversity loss — as the other species contain information on it. On the other hand loosing a species when the number is close to 1 results in the loss of a unique entity.

The pESS approach also forces one to define biodiversity in terms of a specific trait — the one described by the stochastic process. Using a particular trait has the advantage of precision — biodiversity is expressed by the variability of specific entities directly linked to species. In a sense, the pESS links the concept of a species as both a pattern and process [Lidén and Oxelman, 1989]. The process is the evolving trait, an entity that can be directly observed and measured. The patterns are the pre-identified entities on the phylogenetic tree. On the other hand it has the disadvantage of being specific — one looks only at one (or a couple if it is a suite of traits) dimension of the species.

Quantifying the number of species by the pESS of a clade has the advantage of being objective and not subject to potentially arbitrary calls. Not splitting a group is compensated by intra–species variability which can be accommodated by the pESS concept. The need to identify exceptional lineages and possibly novel traits associated with them is discussed by Beaulieu and O’Meara [2016], in the context of clade specific increased/decreased speciation rates. The phylogenetic effective sample size allows for direct comparison between clades with respect to traits, e.g. ones suspected/known of contributing to speciation. Outlier values of pESSs will indicate “interesting” groups of species. Such a methodology combines data from multiple sources, morphological (the traits) and genetic (the phylogeny) — a truly multi-omics approach. With the availability of more and more data from diverse sources mathematical methods that integrate them are being developed more and more in the evolutionary biology world [e.g. Bartoszek and Lió, 2014, Solís-Lemus et al., 2014].

Martins and Hansen [1996] point out that one should expect comparative data sets to contain phylogenetic correlations. It is their absence that should be proved. To prove dependence or independence is a difficult problem in general. One way would be to use information criteria, but it is not clear how many degrees of freedom does the tree have. The relative pESS is an alternative way of showing that phylogenetic correlations are not important. If the value of the relative pESS is close to 1, then the data set is essentially independent.

Maddison and FitzJohn [2015] regret the lack of a method to quantify the number of pseudoreplicates in a phylogenetically correlated dataset. They point out, that the case of discrete traits is even more complicated, as it is the unobserved number of independent origins that matters. Power and p–values, unless one derives model specific tests or uses simulation methods, of e.g. association tests should depend on this number and not on the observed number of species. However, as this number is unknown there is “no quantitative correction to apply to these methods” [Maddison and FitzJohn, 2015], e.g. a contingency table test. The concept of the pESS is what Maddison and FitzJohn [2015] seem to be looking for, but I considered it here in the continuous trait case. Further work is needed to carry the ideas over to the discrete case. However, there is a potential heuristic way of applying the pESS to categorical traits. If one is able to identify continuous traits, that are reasonably related to the discrete one and their pESSs are similar, then their average can be used, as a plug-in for the pESS of the discrete trait in a further downstream analysis/test i.e. an estimator of the number of shifts. The fact that these pESSs are correlated, the traits are dependent through the categorical one and probably between themselves, is actually an advantage. We want the pESS to be nearly the same for each trait and their similarity would indicate sensibility of the described “proxy” approach. If the pESSs for the different traits are dissimilar, then this indicates the need for further investigation, especially choice of traits. The described approach is of course only a suggestion for dealing with discrete, evolutionary correlated data. Further study is needed alongside the development of models where continuous and categorical traits can jointly co-evolve. Another alternative approach to develop in the discrete case, as already mentioned, is the phylogenetic informativeness [Mulder and Crawford, 2015, Townsend, 2007].

The phylogenetic ESS definitions are also interesting from a statistical point of view. The mESS measures the amount of information on the mean value and hence often results in a small pESS, especially in the BM case, where there is limited information on the ancestral state. From all the simulations presented, it seems that the regression ESS captures the amount of independent observations in the data for BM and OU evolution. The good behaviour of the rESS is not surprising as, by construction, it adds up the variance of the independent residuals. Both of these definitions can be used for non–normal processes but we should not expect the regression ESS to be so effective. Rather it would only measure the amount of linearly independent observations. In a general case, I suggest the mutual information ESS, but here work still needs to be done on defining an appropriate *e*(·) transformation in order for 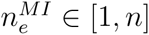 to be in agreement with 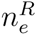 for normal samples.

It could be possible that the proposed pESS approach a step in solving a problem indicated by Faye et al. [2015]: “Unfortunately, not a single of these metrics (providing isolation scores for species - KB) has a strong empirical connection to things we might actually value about biodiversity — trait diversity or trait rarity, evolutionary potential, improved ecosystem function and/or overall genetic information.” The phylogenetic effective sample size forces one to work with a specific trait — if that trait is interesting for biodiversity, then we could have an index that is interesting from Faye et al. [2015] point of view. What is more important, pESSs are cheap to obtain.

## Acknowledgments

I was supported by the Knut and Alice Wallenberg Foundation. I would like to like to acknowledge two anonymous reviewers, whose comments greatly improved the manuscript. I would like to thank Chad Eliason for providing me with the ducks data and Marcin Piwczyński for the Mantellidae and *Chaerophyllum* data. I would like to thank Bengt Oxelman and Tanja Stadler for encouragement and numerous comments on this work.

## Phylogenetic effective sample size Supplementary histograms

Krzysztof Bartoszek

Department of Mathematics, Uppsala University, 751 06 Uppsala, Sweden.

**Figure S.1:**
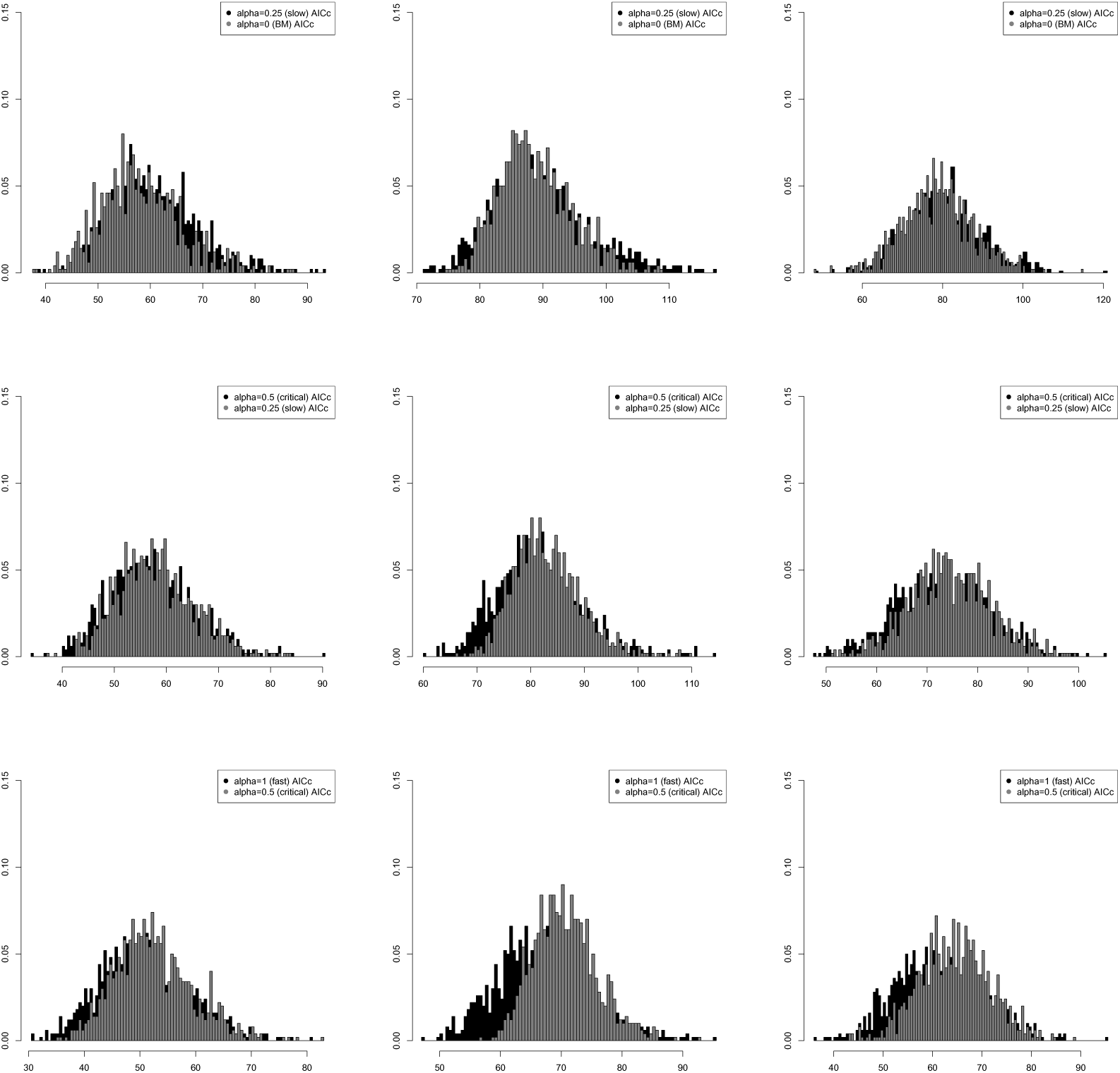
Histograms of AIC_c_ values with 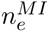 effective sample size correction for different types of trees and evolutionary processes. The sample sizes are *n* = 30 (left unbalanced tree and Yule) and *n* = 32 (balanced tree). First column: balanced tree, second column: left unbalanced tree, third column: 1000 pure–birth Yule trees (*λ* = 1). The balanced trees and unbalanced trees were generated using the function stree() of the R ape package, the Yule trees by the TreeSim R package. First row: Ornstein–Uhlenbeck process (*α* = 0.25, *σ*^2^ = 1, *X*_0_ = 0, *θ* = 0 black true model), Brownian motion (*X*_0_ = 0, *σ*^2^ = 1 gray alternative model), second row: Ornstein–Uhlenbeck process (*α* = 0.5, *σ*^2^ = 1, *X*_0_ = 0, *θ* = 0 black true model), Ornstein–Uhlenbeck process (*α* = 0.25, *σ*^2^ = 1, *X*_0_ = 0, *θ* = 0 gray alternative model), third row: Ornstein–Uhlenbeck process (*α* = 1, *σ*^2^ = 1, *X*_0_ = 0, *θ* = 0 black true model), fourth row: Ornstein–Uhlenbeck process (*α* = 0.5, *σ*^2^ = 1, *X*_0_ = 0, *θ* = 0 gray alternative model). We simulate data under both the true and alternative evolutionary models 1000 times and then calculate AIC_*c*_ values for each simulated pair.

**Figure S.2:**
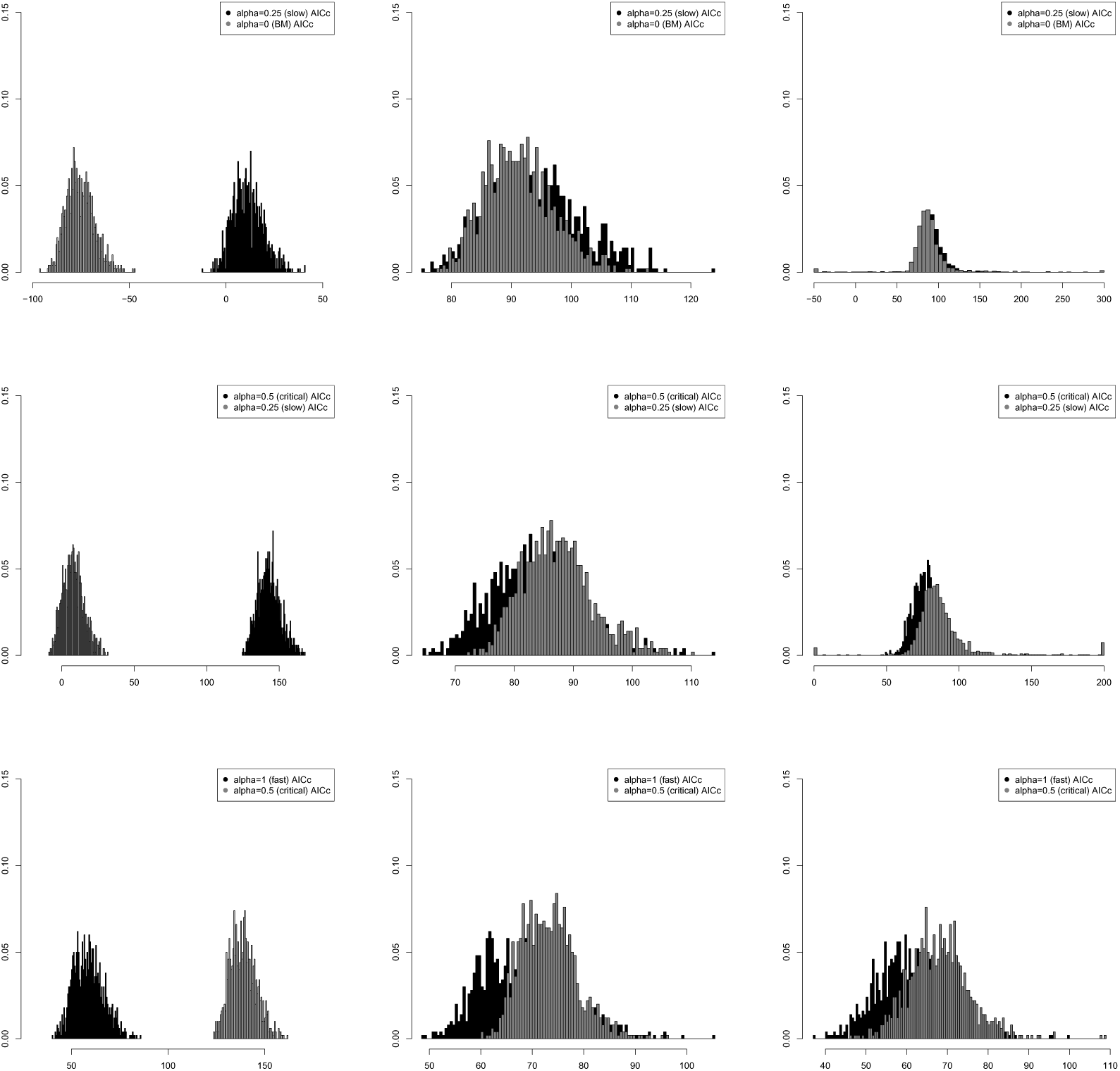
Histograms of AIC_*c*_ values with 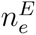 effective sample size correction for different types of trees and evolutionary processes. The sample sizes are *n* = 30 (left unbalanced tree and Yule) and *n* = 32 (balanced tree). First column: balanced tree, second column: left unbalanced tree, third column: 1000 pure–birth Yule trees (*λ* = 1). The balanced trees and unbalanced trees were generated using the function stree() of the R ape package, the Yule trees by the TreeSim R package. First row: Ornstein–Uhlenbeck process (*α* = 0.25, *σ*^2^ = 1, *X*_0_ = 0, *θ* = 0 black true model), Brownian motion (*X*_0_ = 0, *σ*^2^ = 1 gray alternative model), second row: Ornstein–Uhlenbeck process (*α* = 0.5, *σ*^2^ = 1, *X*_0_ = 0, *θ* = 0 black true model), Ornstein–Uhlenbeck process (*α* = 0.25, *σ*^2^ = 1, *X*_0_ = 0, *θ* = 0 gray alternative model), third row: Ornstein–Uhlenbeck process (*α* = 1, *σ*^2^ = 1, *X*_0_ = 0, *θ* = 0 black true model), fourth row: Ornstein–Uhlenbeck process (*α* = 0.5, *σ*^2^ = 1, *X*_0_ = 0, *θ* = 0 gray alternative model). We simulate data under both the true and alternative evolutionary models 1000 times and then calculate AIC_*c*_ values for each simulated pair.

**Figure S.3:**
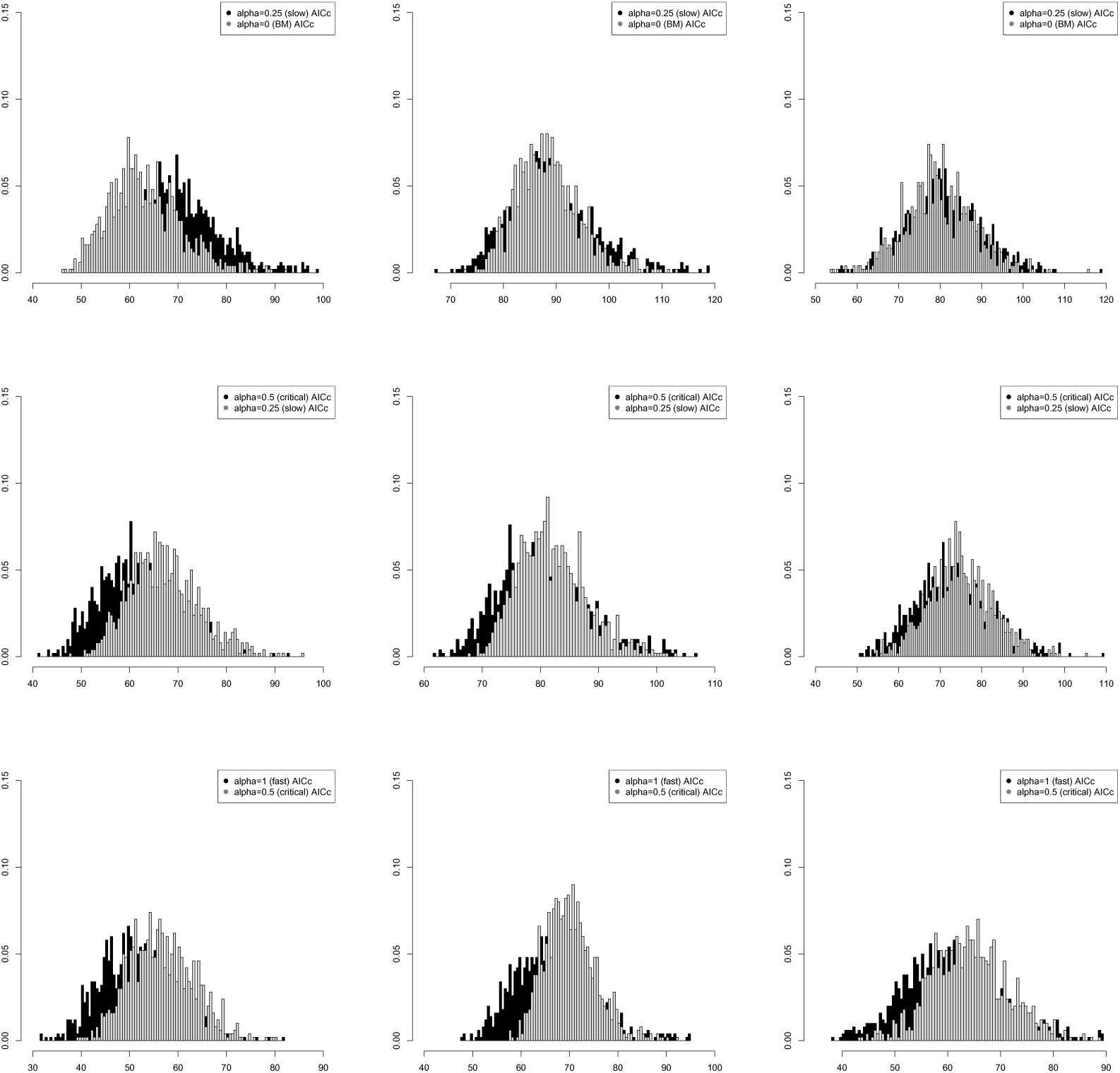
Histograms of AIC_*c*_ values with 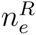 effective sample size correction for different types of trees and evolutionary processes. The sample sizes are *n* = 30 (left unbalanced tree and Yule) and *n* = 32 (balanced tree). First column: balanced tree, second column: left unbalanced tree, third column: 1000 pure–birth Yule trees (*λ* = 1). The balanced trees and unbalanced trees were generated using the function stree() of the R ape package, the Yule trees by the TreeSim R package. First row: Ornstein–Uhlenbeck process (*α* = 0.25, *σ*^2^ = 1, *X*_0_ = 0, *θ* = 0 black true model), Brownian motion (*X*_0_ = 0, *σ*^2^ = 1 gray alternative model), second row: Ornstein–Uhlenbeck process (*α* = 0.5, *σ*^2^ = 1, *X*_0_ = 0, *θ* = 0 black true model), Ornstein–Uhlenbeck process (*α* = 0.25, *σ*^2^ = 1, *X*_0_ = 0, *θ* = 0 gray alternative model), third row: Ornstein–Uhlenbeck process (*α* = 1, *σ*^2^ = 1, *X*_0_ = 0, *θ* = 0 black true model), fourth row: Ornstein–Uhlenbeck process (*α* = 0.5, *σ*^2^ = 1, *X*_0_ = 0, *θ* = 0 gray alternative model). We simulate data under both the true and alternative evolutionary models 1000 times and then calculate AIC_*c*_ values for each simulated pair.

**Figure S.4:**
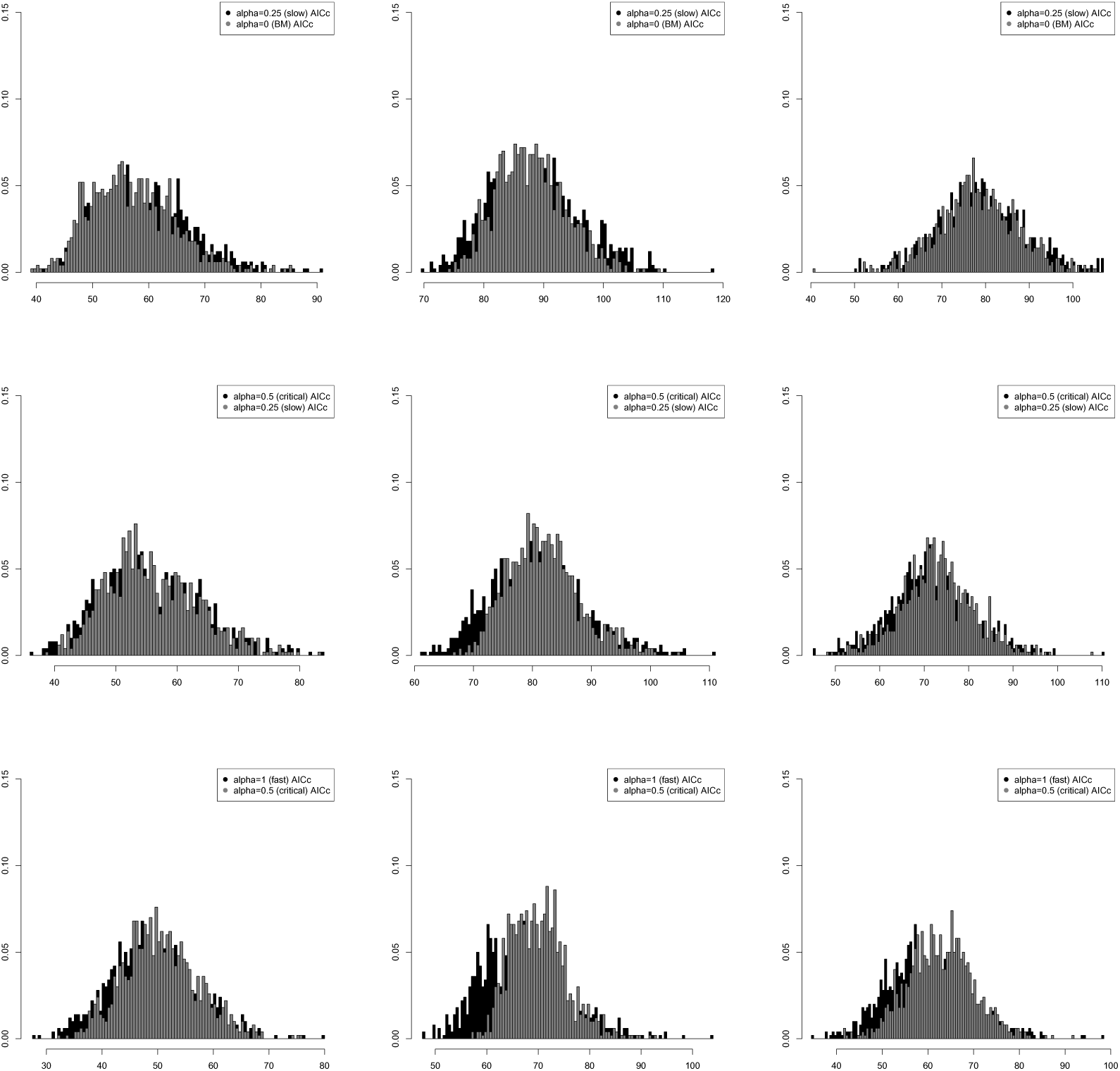
Histograms of AIC_c_ values with no effective sample size correction for different types of trees and evolutionary processes. The sample sizes are *n* = 30 (left unbalanced tree and Yule) and *n* = 32 (balanced tree). First column: balanced tree, second column: left unbalanced tree, third column: 1000 pure–birth Yule trees (*λ* = 1). The balanced trees and unbalanced trees were generated using the function stree() of the R ape package, the Yule trees by the TreeSim R package. First row: Ornstein–Uhlenbeck process (*α* = 0.25, *σ*^2^ = 1, *X*_0_ = 0, *θ* = 0 black true model), Brownian motion (*X*_0_ = 0, *σ*^2^ = 1 gray alternative model), second row: Ornstein–Uhlenbeck process (*α* = 0.5, *σ*^2^ = 1, *X*_0_ = 0, *θ* = 0 black true model), Ornstein–Uhlenbeck process (*α* = 0.25, *σ*^2^ = 1, *X*_0_ = 0, *θ* = 0 gray alternative model), third row: Ornstein–Uhlenbeck process (*α* = 1, *σ*^2^ = 1, *X*_0_ = 0, *θ* = 0 black true model), fourth row: Ornstein–Uhlenbeck process (*α* = 0.5, *σ*^2^ = 1, *X*_0_ = 0, *θ* = 0 gray alternative model). We simulate data under both the true and alternative evolutionary models 1000 times and then calculate AIC_c_ values for each simulated pair.

**Figure S.5:**
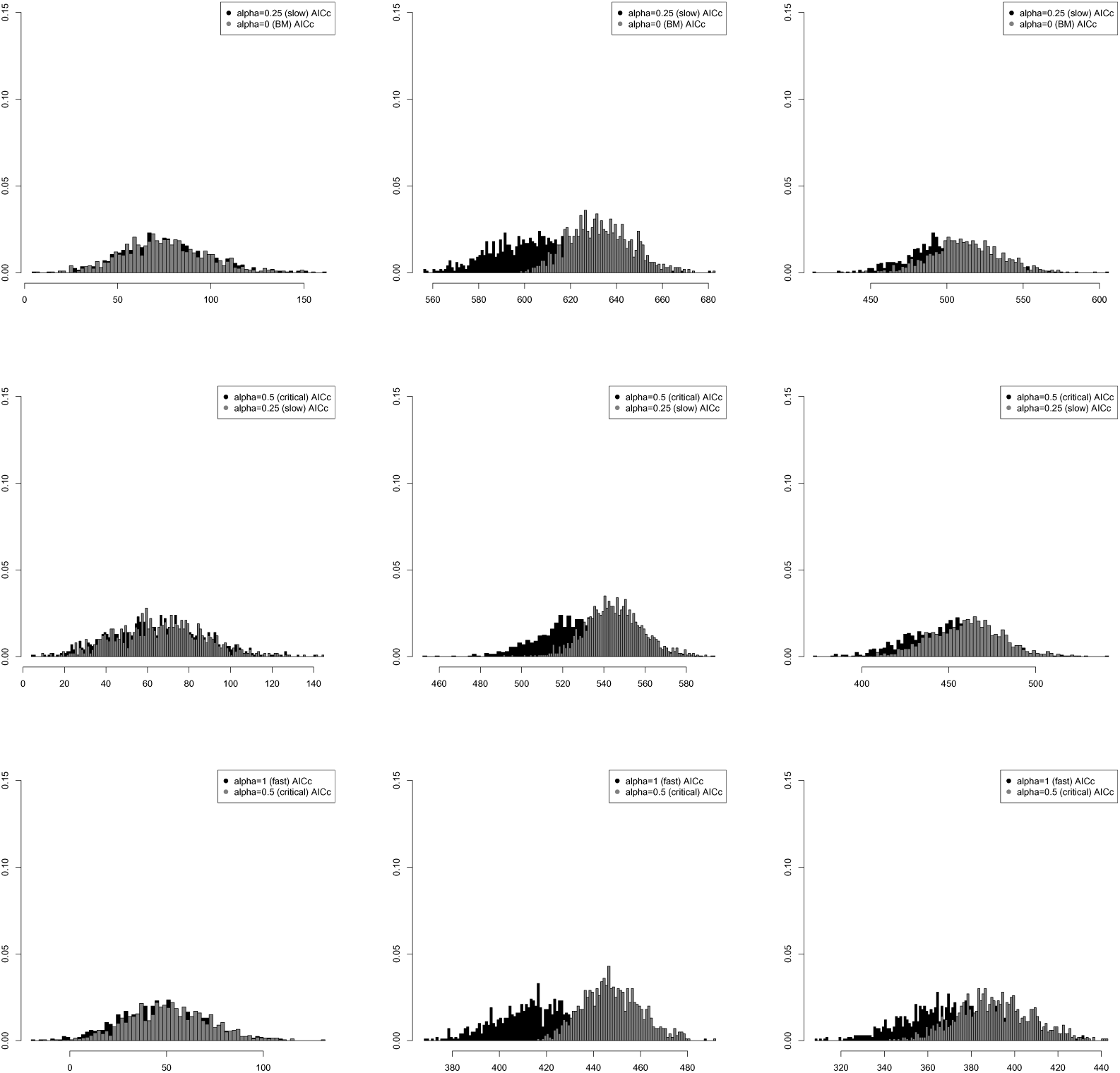
Histograms of AIC_*c*_ values with 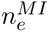 effective sample size correction for different types of trees and evolutionary processes. The sample sizes are *n* = 205 (left unbalanced tree and Yule) and *n* = 256 (balanced tree). First column: balanced tree, second column: left unbalanced tree, third column: 1000 pure–birth Yule trees (*λ* = 1). The balanced trees and unbalanced trees were generated using the function stree() of the R ape package, the Yule trees by the TreeSim R package. First row: Ornstein–Uhlenbeck process (*α* = 0.25, *σ*^2^ = 1, *X*_0_ = 0, *θ* = 0 black true model), Brownian motion (*X*_0_ = 0, *σ*^2^ = 1 gray alternative model), second row: Ornstein–Uhlenbeck process (*α* = 0.5, *σ*^2^ = 1, *X*_0_ = 0, *θ* = 0 black true model), Ornstein–Uhlenbeck process (*α* = 0.25, *σ*^2^ = 1, *X*_0_ = 0, *θ* = 0 gray alternative model), third row: Ornstein–Uhlenbeck process (*α* = 1, *σ*^2^ = 1, *X*_0_ = 0, *θ* = 0 black true model), fourth row: Ornstein–Uhlenbeck process (*α* = 0.5, *σ*^2^ = 1, *X*_0_ = 0, *θ* = 0 gray alternative model). We simulate data under both the true and alternative evolutionary models 1000 times and then calculate AIC_*c*_ values for each simulated pair.

**Figure S.6:**
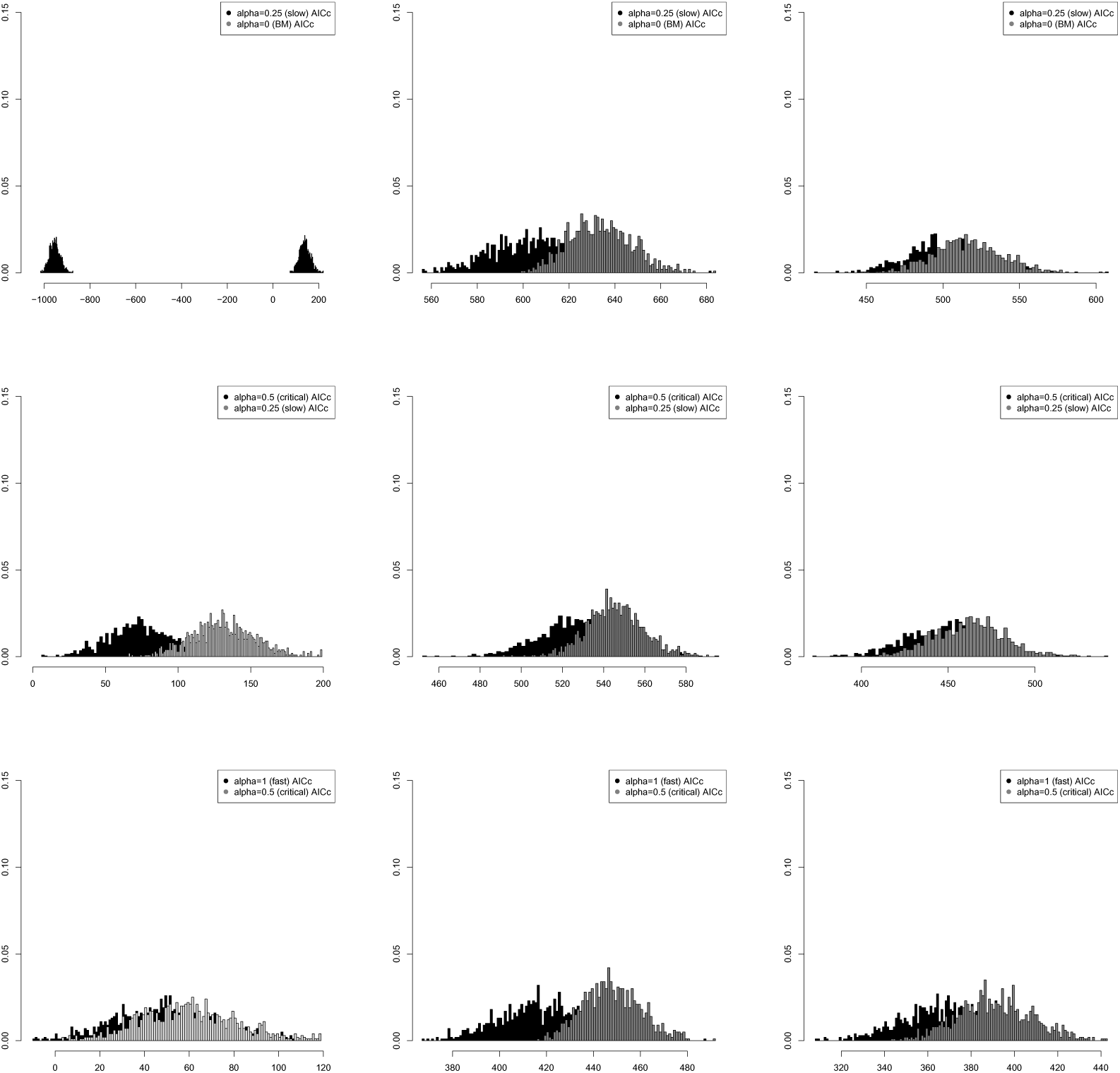
Histograms of AIC_*c*_ values with 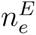 effective sample size correction for different types of trees and evolutionary processes. The sample sizes are *n* = 205 (left unbalanced tree and Yule) and *n* = 256 (balanced tree). First column: balanced tree, second column: left unbalanced tree, third column: 1000 pure–birth Yule trees (*λ* = 1). The balanced trees and unbalanced trees were generated using the function stree() of the R ape package, the Yule trees by the TreeSim R package. First row: Ornstein–Uhlenbeck process (*α* = 0.25, *σ*^2^ = 1, *X*_0_ = 0, *θ* = 0 black true model), Brownian motion (*X*_0_ = 0, *σ*^2^ = 1 gray alternative model), second row: Ornstein–Uhlenbeck process (*α* = 0.5, *σ*^2^ = 1, *X*_0_ = 0, *θ* = 0 black true model), Ornstein–Uhlenbeck process (*α* = 0.25, *σ*^2^ = 1, *X*_0_ = 0, *θ* = 0 gray alternative model), third row: Ornstein–Uhlenbeck process (*α* = 1, *σ*^2^ = 1, *X*_0_ = 0, *θ* = 0 black true model), fourth row: Ornstein–Uhlenbeck process (*α* = 0.5, *σ*^2^ = 1, *X*_0_ = 0, *θ* = 0 gray alternative model). We simulate data under both the true and alternative evolutionary models 1000 times and then calculate AIC_*c*_ values for each simulated pair.

**Figure S.7:**
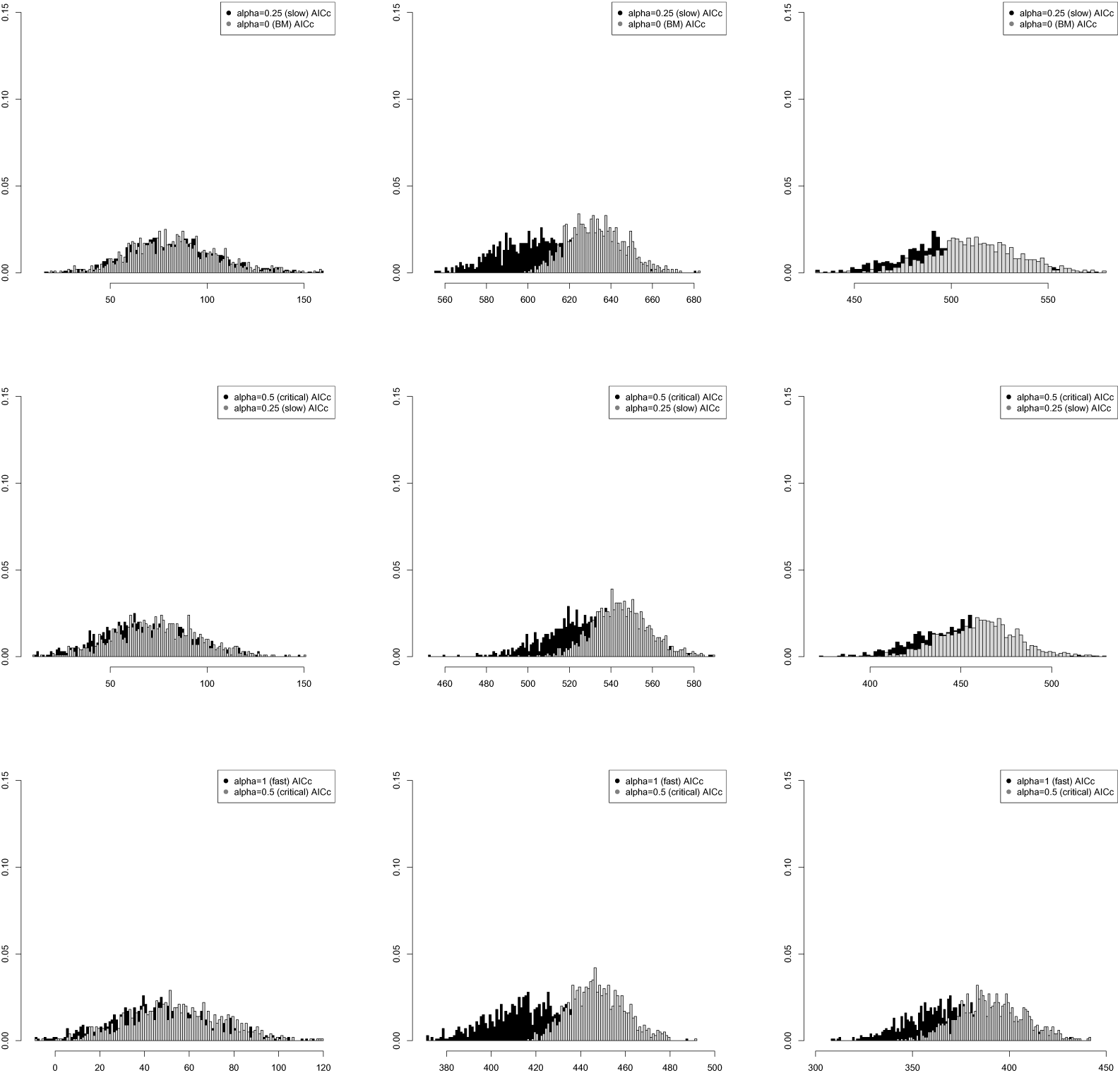
Histograms of AIC_*c*_ values with 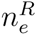 effective sample size correction for different types of trees and evolutionary processes. The sample sizes are *n* = 205 (left unbalanced tree and Yule) and *n* = 256 (balanced tree). First column: balanced tree, second column: left unbalanced tree, third column: 1000 pure–birth Yule trees (*λ* = 1). The balanced trees and unbalanced trees were generated using the function stree() of the R ape package, the Yule trees by the TreeSim R package. First row: Ornstein–Uhlenbeck process (*α* = 0.25, *σ*^2^ = 1, *X*_0_ = 0, *θ* = 0 black true model), Brownian motion (*X*_0_ = 0, *σ*^2^ = 1 gray alternative model), second row: Ornstein–Uhlenbeck process (*α* = 0.5, *σ*^2^ = 1, *X*_0_ = 0, *θ* = 0 black true model), Ornstein–Uhlenbeck process (*α* = 0.25, *σ*^2^ = 1, *X*_0_ = 0, *θ* = 0 gray alternative model), third row: Ornstein–Uhlenbeck process (*α* = 1, *σ*^2^ = 1, *X*_0_ = 0, *θ* = 0 black true model), fourth row: Ornstein–Uhlenbeck process (*α* = 0.5, *σ*^2^ = 1, *X*_0_ = 0, *θ* = 0 gray alternative model). We simulate data under both the true and alternative evolutionary models 1000 times and then calculate AIC_*c*_ values for each simulated pair.

**Figure S.8:**
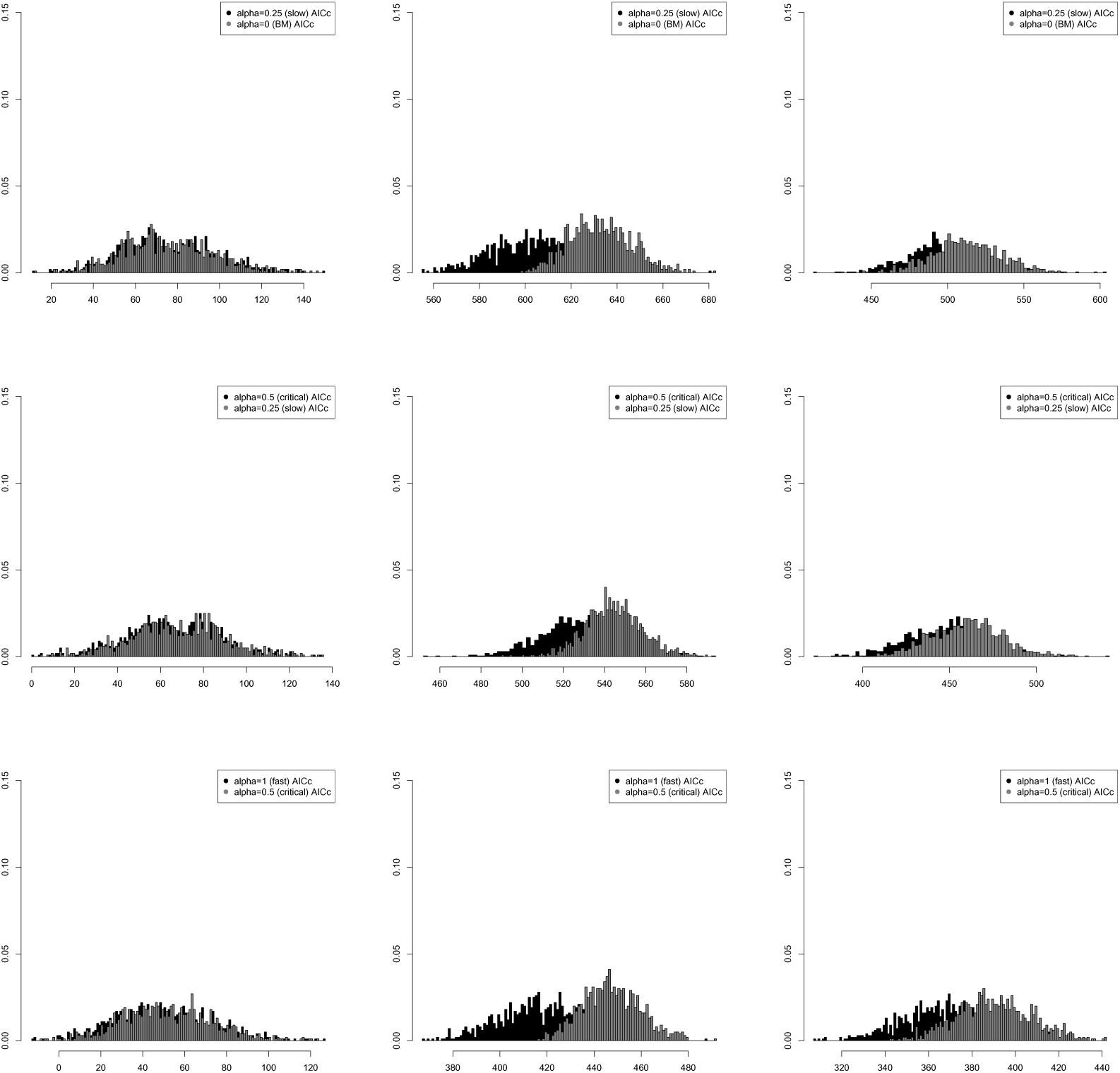
Histograms of AIC_*c*_ values with no effective sample size correction for different types of trees and evolutionary processes. The sample sizes are *n* = 205 (left unbalanced tree and Yule) and *n* = 256 (balanced tree). First column: balanced tree, second column: left unbalanced tree, third column: 1000 pure–birth Yule trees (*λ* = 1). The balanced trees and unbalanced trees were generated using the function stree() of the R ape package, the Yule trees by the TreeSim R package. First row: Ornstein–Uhlenbeck process (*α* = 0.25, *σ*^2^ = 1, *X*_0_ = 0, *θ* = 0 black true model), Brownian motion (*X*_0_ = 0, *σ*^2^ = 1 gray alternative model), second row: Ornstein–Uhlenbeck process (*α* = 0.5, *σ*^2^ = 1, *X*_0_ = 0, *θ* = 0 black true model), Ornstein–Uhlenbeck process (*α* = 0.25, *σ*^2^ = 1, *X*_0_ = 0, *θ* = 0 gray alternative model), third row: Ornstein–Uhlenbeck process (*α* = 1, *σ*^2^ = 1, *X*_0_ = 0, *θ* = 0 black true model), fourth row: Ornstein–Uhlenbeck process (*α* = 0.5, *σ*^2^ = 1, *X*_0_ = 0, *θ* = 0 gray alternative model). We simulate data under both the true and alternative evolutionary models 1000 times and then calculate AIC_*c*_ values for each simulated pair.

